# GFHunter enables accurate and efficient gene fusion detection in long-read cancer transcriptomes

**DOI:** 10.1101/2025.02.23.639788

**Authors:** Yadong Liu, Zhenhao Lu, Yadong Wang, Tao Jiang

## Abstract

The precise identification of gene fusions is crucial for cancer diagnosis and therapeutic decision-making. Long-read transcriptome sequencing provides distinct advantages over short-read technologies by capturing full-length fusion gene structures. However, fully harnessing long-read data for cancer research necessitates advanced computational approaches. In this study, we present GFHunter, a novel computational framework designed for efficient and accurate gene fusion detection. Benchmarking on both simulated and real long-read transcriptome datasets from non-tumor and cancer cell lines demonstrates that GFHunter accurately detects gene fusions with high sensitivity and significantly reduces false positives. Additionally, GFHunter runs 2-3 times faster and requires only 16%-50% of the memory compared to state-of-the-art tools. Notably, GFHunter uniquely identifies two known cancer-related fusions in HCT-116 and SKBR-3 cancer cell lines. These results highlight GFHunter’s potential as a powerful tool for advancing precision oncology and molecular diagnostics.

## Introduction

A series biological mechanisms and processes such as chromosomal rearrangements or trans-splicing leading to the formation of gene fusions^1–3^. Gene fusion typically encodes abnormal fusion proteins, which may diff in structure and function from the normal proteins, that resulting the changes in cellular functions, such as aberrant proliferation, survival, or migration, and is frequent drivers in certain cancer types^1, 4, 5^. The Cancer Genome Atlas (TCGA) estimates that gene fusion drove 16.5% of human cancers by comprehensive analysis across a range of tumor types^6^. For example, the *BCR-ABL1* fusion was observed in chronic myeloid leukemia^7^; *TMPRSS2-ERG* fusion was found in ∼50% of prostate cancer^8^; *DNAJB1-PRKACA* fusion was considered to be the hallmark and likely driver of fibrolamellar carcinoma^9^; moreover, gene fusion is also important for other diseases^10^. Thus, gene fusion can be used as potential biomarkers and therapeutic targets for cancer diagnosis in breast cancer^11^, ovarian cancer^12^, prostate cancer^13^ and others. Recently years, the U.S. Food and Drug Administration and the National Medical Products Administration of China has accelerated approval drugs for the treatment of cancers harboring gene fusions^14^. Hence, better understanding gene fusions is a routine task in genomics-guided precision oncology and lead to the development of novel targeted therapies in the future.

Over the past decade, short-read transcriptome sequencing has emerged as a powerful strategy for capturing the landscape of gene fusions in cancer, playing a pivotal role in advancing precision medicine^15–17^. To this end, a wide array of tools and pipelines have been developed, broadly categorized into alignment-based and assembly-based approaches. Alignment-based tools, such as STAR-Fusion^18^, Arriba^19^, deFuse^20^, SOAPfuse^21^ and TopHat-Fusion^22^, identify gene fusions by aligning short reads to a reference genome (or transcriptome) and subsequently analyzing discordant alignment signatures. In contrast, assembly-based tools, including JAFFA^23^, FuSeq^24^ and nFuse^25^, assembly short reads into longer transcripts followed by identification of chimeric transcripts and corresponding breakpoints relevant to chromosomal rearrangements. Despite their utility, several assessment studies^19, 26, 27^ have highlighted significant challenges in accurately identifying chimeric reads or discordant read pairs and detecting precise gene fusions from short-read data. These challenges stem from the inherent limitations of short-read sequencing, such as restricted read lengths (typically ranging from 50 to 300 base pairs), which can lead to inaccuracies in determining supporting reads for gene fusion events, particularly in repetitive or low-complexity genomic regions. Additionally, the complex splicing structures of isoforms further complicate the accurate identification of gene fusions.

Long-read sequencing technologies, including Pacific Bioscience (PacBio)^28^ and Oxford Nanopore Technologies (ONT)^29^ can generate long reads up to several mega-bases, which offers an unprecedented opportunity to capture full length transcripts^30, 31^. The long-range information may alleviate the challenges from short reads and provide promising ability to detect gene fusions, which can improve our understanding transcriptomic complexity and present new perspectives for gene fusion in cancer initiation and progression. Both PacBio and ONT support cDNA sequencing, and ONT additionally offers direct RNA (dRNA) sequencing^28, 29^, which allows for the direct observation of full-length transcripts without the need for cDNA conversion. While the use of full-length transcripts for gene fusion detection may seem straightforward, it presents several non-trivial challenges. These include: 1) distinguishing true fusion events from confounding biological phenomena such as exon skipping or alternative splicing; 2) the potential loss of genuine fusion signals due to misalignment artifacts or the presence of highly homologous isoforms, leading to signal attenuation or complete omission; and 3) the inherent difficulty in precisely resolving fusion junctions, exacerbated by the elevated error rates of long-read transcriptome sequencing and systemic biases in alignment scoring algorithms. To address these challenges, several computational tools have been proposed for long-read RNA-Seq data, such as Genion^32^, LongGF^33^, JAFFAL^34^ and FusionSeeker^35^.

With the exception of JAFFAL, most tools align long transcriptome reads to the reference genome to detect gene fusions. For instance, Genion identifies chains of exons, clusters reads to define and rank fusion candidates, and ultimately determines true fusion events using a statistical approach. LongGF leverages multiple mapped long reads to identify candidate gene pairs, filters for gene pairs with non-random supporting reads, and outputs a prioritized list of candidate gene fusions. FusionSeeker scans read alignments for candidate fusions and applies density-based spatial clustering (DBSCAN) to generate fusion calls. Additionally, FusionSeeker employs several filtration steps and performs multiple sequence alignment to generate a consensus transcript sequence. In contrast, JAFFAL is the only one tool that align reads into transcriptome, it is built on the concepts developed in JAFFA^23^ and overcomes the high error rate in long reads with several filtering heuristics. From my perspective, aligning long reads to the reference transcriptome offers several advantages, such as 1) it eliminates the need to identify introns, thereby improving alignment accuracy; 2) it is computationally efficient and requires fewer resources, as reads are aligned only to transcribed regions, which constitute approximately 3% of the entire genome^36^; and 3) it enables more reads are involved in gene fusion detection, as all reads are aligned to the gene regions.

Existing methods still face limitations in achieving both high sensitivity and accuracy, particularly when it comes to pinpointing breakpoints with precision and generating accurate fusion transcripts. Hence, we introduce GFHunter, a novel tool designed for the accurate and efficient identification of gene fusions, along with the precise reconstruction of fused sequences, using long transcriptome sequencing data. GFHunter demonstrates superior performance compared to state-of-the-art methods, as validated through comprehensive benchmarking on both simulated data and real-world cancer cell line data from Oxford Nanopore Technologies (ONT) cDNA, direct RNA (dRNA), and PacBio Iso-seq sequencing. The exceptional performance mainly derived from four key points innovations, 1) GFHunter rebuilds the reference transcriptome by retaining only protein-coding transcripts, enabling efficient alignment of long reads to the transcriptome; 2) GFHunter employs sophisticated filters to identify signal reads and utilizes sparse matrix multiplication to pair candidate fusion genes; 3) GFHunter clusters fusion breakpoints though a bottom-up hierarchical clustering, and performs multiple sequence alignment (abPOA^37^) to generate accurate fusion transcript sequence; and 4) GFHunter realign the fused transcript sequence to reference genome to refine the breakpoints and rank fusions.

## Results

### Overview of GFHunter approach

GFHunter is an accurate and efficient gene fusion detection tool designed for long-read transcriptome sequencing data, utilizing a transcriptome-based alignment approach. GFHunter comprises three major steps, each tailored to the structure of the fusion genes (*Figure 1a*) and alignment signatures to achieve highly sensitive and precise gene fusion calls (Online Methods).

**Figure 1.**
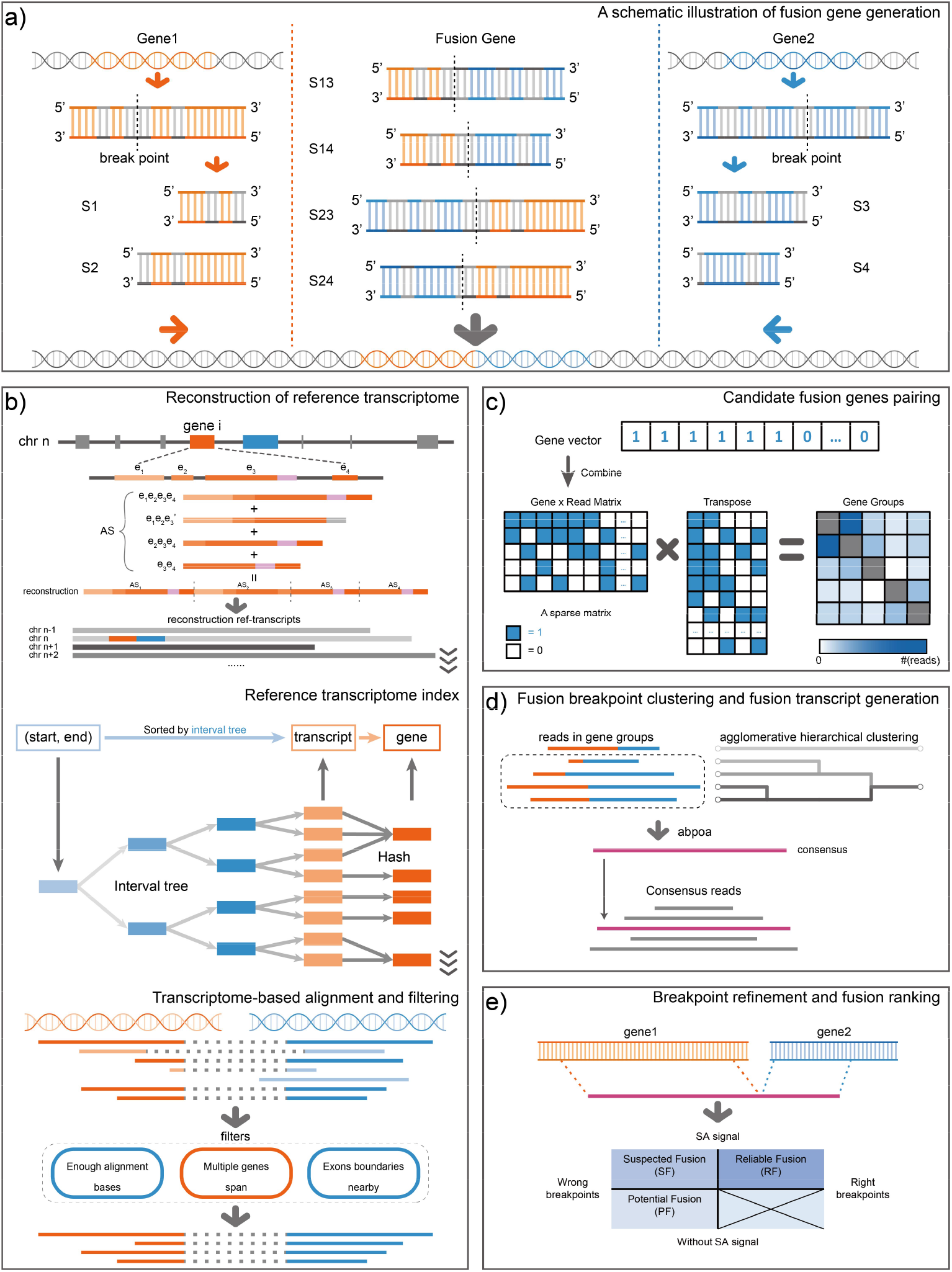
A schematic illustration of GFHunter. **a**. Mechanism of fusion gene generation. Gene 1 and Gene 2 can form four potential fusion types: S13, S14, S23, or S24. **b**. Transcriptome-based long-read alignment. Firstly, GFHunter reconstructs the reference transcriptome by concatenating all the selected transcript sequences. Subsequently, it constructs a hash-based transcriptome index via an interval tree to map genes according to their genomic positions. Lastly, reads are aligned to the reconstructed transcriptome using Minimap2, followed by filtering to preserve potential fusion supporting reads. **c**. Candidate fusion gene pairing. GFHunter identifies potential fusion pairs through sparse matrix multiplication, where “1” indicates a read spans the gene and “0” otherwise. **d**. Fusion breakpoints clustering and transcript generation. GFHunter clusters reads (breakpoints) though bottom-up hierarchical clustering strategy, and applies multiple sequence alignment to generate fusion transcripts. **e**. Breakpoint refinement and fusion ranking. GFHunter refines breakpoints and ranks fusion events by cross-referencing results from both the transcriptome and genome.

Step1: GFHunter begins by reconstructing the reference transcriptome, focusing primarily on protein-coding genes. A hash-based index is employed to efficiently retrieve specific genes via an interval tree. Subsequently, GFHunter performs transcriptome-based alignment by mapping long RNA-seq reads to the reconstructed reference transcriptome. This approach minimizes the challenges associated with frequent splicing events and reduces ambiguous or heterogeneous alignments. To further enhance precision, several sophisticated filters are applied to retain potential gene fusion signatures, thereby improving the accuracy and efficiency of downstream fusion detection (*Figure 1b*).

Step2: GFHunter represents the alignments within genes as a sparse 0-1 matrix, and utilizes sparse matrix multiplication between the matrix and its transpose to generate a symmetric matrix. This matrix captures the co-occurrence patterns of reads spanning different gene pairs, enabling efficient identification of plausible gene fusions. Initial fusion breakpoints are determined based on alignment flags and the relative positions of the genes, allowing precise localization of whether breakpoints occur before or after specific exons (*Figure 1c*).

Step3: GFHunter organizes inferred fusion breakpoint pair into 2D coordinate point, which are then subjected to bottom-up hierarchical clustering based on Manhattan distances. Clusters with distances shorter than a predefined cutoff are merged. For each merged cluster, local sequences between fusion breakpoint pairs are extracted, and multiple sequence alignment is performed using abPOA to generate precise fusion transcript sequences (*Figure 1d*).

Step4: GFHunter performs realignment of the fusion transcript to the reference genome to reconstruct the fusion event, leveraging long-range alignment and precise splicing site identification (genome-based alignment). Subsequently, GFHunter refines and ranks the confidence level of each fusion event by cross-referencing the results from both the reference transcriptome and genome. This process evaluates three distinct scenarios to comprehensively assess the reliability of each identified fusion event (*Figure 1e*).

GFHunter ranks gene fusions into three distinct confidence levels based on the concordance between genome-based and transcriptome-based fusion breakpoints: 1) “Reliable” fusions, the highest-confidence category includes gene fusions that are strongly supported by genome-based spliced alignment of the fusion transcript, exhibiting correct split alignment signals. 2) “Suspected” fusions, the intermediate category includes fusions where the inferred breakpoints from hierarchical clustering show inconsistency with the spliced alignment of the fusion transcript, despite having sufficient supporting reads and belonging to the same gene pair. 3) “Potential” fusions, the lowest-confidence category includes fusions where the fusion transcript cannot be aligned to two distinct genes. The definition of different ranks refers to the *Online Method*. By categorizing fusions into these three levels, GFHunter provides a nuanced confidence assessment, enabling researchers to prioritize high-confidence fusions for downstream analysis while flagging lower-confidence events for further validation.

Eventually, GFHunter reports each gene fusion event with the corresponding gene pairs, fusion breakpoints within each gene, rank class, supporting reads information, and the fusion transcript. We conducted a comprehensive evaluation of GFHunter with four integral competing approaches, i.e., Genion, LongGF, JAFFAL and FusionSeeker, using diverse datasets from real-world well-studied cancer samples and specially designed simulations, to assess accuracy and robustness of gene fusion event identification across different scenarios.

### Fusion identification on simulated datasets

We first assess the accuracy of gene fusion detection on positive and negative simulations. The positive simulation, with fusion transcripts, and the negative simulation, without fusion transcripts, were generated from GENCODE gene annotations (release 44 used in this study, optional). For positive simulation, fusion transcripts were specifically designed by selecting gene pairs from both inter-chromosomal and intra-chromosomal regions, with breakpoints randomly assigned (*Figure 2a*, details refer to *Supplementary Notes*). A total of 500 fusion transcripts were reconstructed, with 470 being inter-chromosomal and 30 intra-chromosomal. *Figure 2b* illustrates the distribution of gene pairs across chromosomes, revealing a positive correlation between the number of fusion genes and the gene counts on each chromosome, which underscores the randomness of fusion events (*Supplementary Table 1*). For negative simulation, 249,443 normal transcripts from annotations are adopted to imitate non-tumor cell line transcriptome. The length distributions of the normal transcripts and fusion transcripts are shown in *Figure 2c* and *Figure 2d*, with N50 lengths of 687 and 2574 base pairs, respectively. Subsequently, two integral platforms, that is PacBio Iso-seq-like and ONT cDNA-like reads from the transcript sequences were generated using pbsim3^38^, with 4 different sequencing coverages, i.e., 10×, 20×, 30×, 50×.

**Table 1.** Reported fusions on real non-tumor datasets (HG002)

| HG002 | Reported fusions of each tool |  |  |  |  |  |  | Fusions supported by number of tools |  |  |  |  |
| --- | --- | --- | --- | --- | --- | --- | --- | --- | --- | --- | --- | --- |
|  | GFHunter |  | LongGF | JAFFAL |  | FusionSeeker | Genion | 1 | 2 | 3 | 4 | 5 |
|  | RF | RF+SF |  | HC | HC+LC |  |  |  |  |  |  |  |
| cDNA | 2 | 9 | 42 | 3 | 24 | 383 | 24 | 386 | 34 | 0 | 0 | 0 |
| dRNA | 2 | 5 | 5 | 8 | 9 | 36 | 34 | 75 | 3 | 0 | 1 | 0 |
| Iso-seq | 8 | 16 | 128 | 16 | 38 | 45 | 98 | 263 | 14 | 0 | 1 | 0 |

RF: Reliable fusions; SF: Suspected fusions; HC: High-confidence; LC: Low-confidence

**Figure 2.**
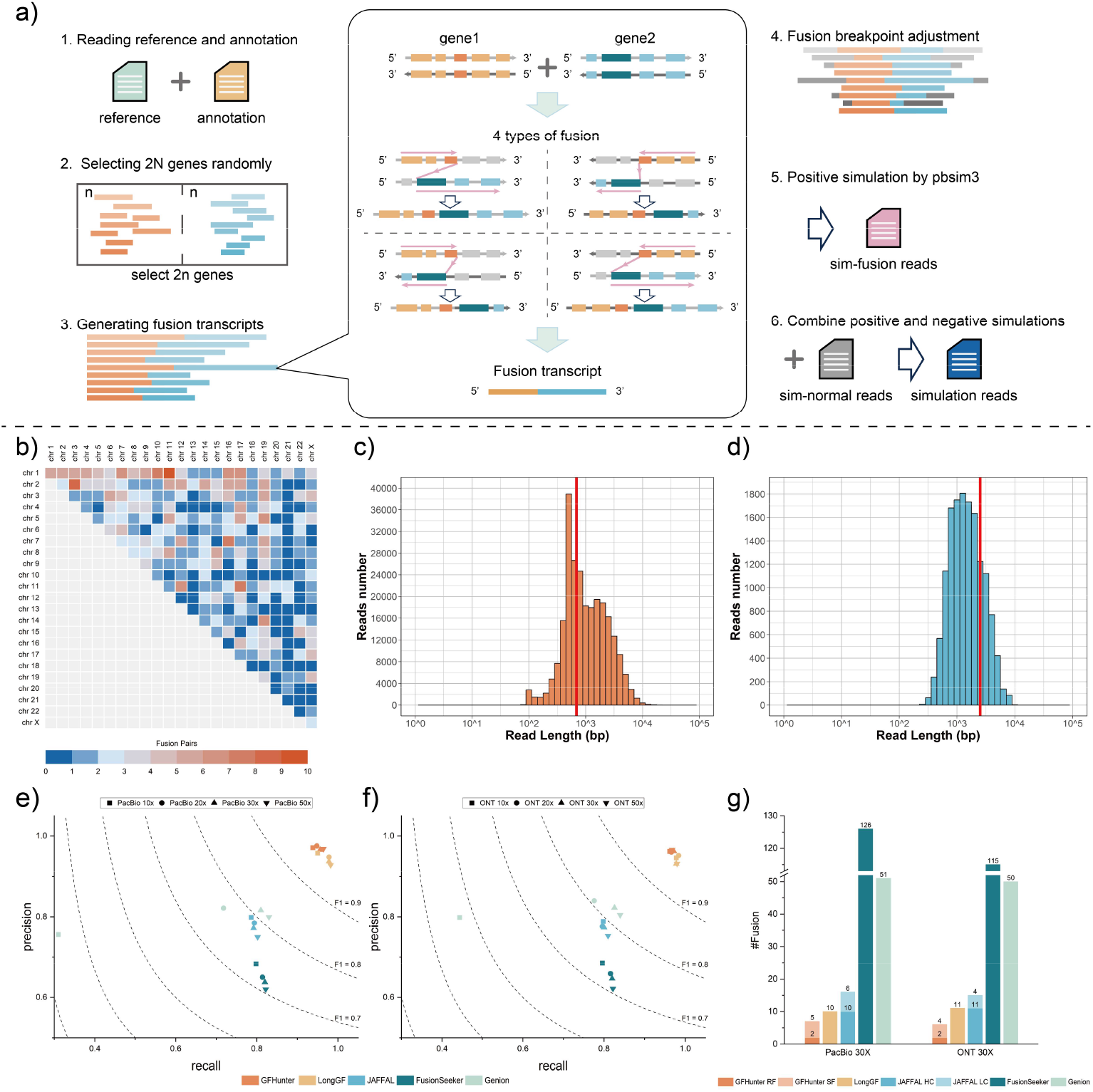
Benchmarking results on simulation datasets. **a**. Schematic representation of the simulation workflow. **b**. Heatmap showing the distribution of simulated fusion genes across chromosomes (chr1 to chrX). **c**. N50 distribution for Negative simulation. **c&d**. Transcript lengths distribution for negative(c) and positive (d) simulations. The red line indicates the N50 length. **e&f**. Contour map of recall, precision and F1-score for PacBio (e) and ONT (f) datasets at varying sequencing coverages (10×, 20×, 30×, 50×). **g**. Number of detected fusions on negative simulated datasets (PacBio 30×, ONT 30×).

We combine the positive and negative simulations into 8 individual synthetic datasets (2 platforms and 4 coverages). For benchmarking, we considered “Reliable” and “Suspected” fusions for GFHunter, and “High Confidence” and “Low Confidence” fusions for JAFFAL. Across all datasets, GFHunter and LongGF achieved the higher yields, significantly outperformed other competing methods, demonstrating their superior ability and versatility in gene fusion identification (*Supplementary Table 2*). Moreover, GFHunter exhibited remarkable accuracy, surpassing LongGF in precision by 1.3%, 2.7%, 2.9%, 4.0%, 1.6%, 0.8%, 3.1%, and 3.4% across the eight datasets, respectively. Overall, GFHunter achieved F1 scores exceeding 95% (95.4%, 96.1%, 96.1%, 96.6% for PacBio, and 96.2%, 96.3%, 96.7%, 96.7% for ONT at 10×, 20×, 30×, 50×, respectively). LongGF achieved similar F1 scores but was slightly lower, with an average reduction of 0.44%. In contrast, the F1 scores of JAFFAL, FusionSeeker, and Genion consistently hovered around 80% or lower, highlighting their limited ability to accurately detect gene fusions.

Under different sequencing coverage, GFHunter and LongGF demonstrated stable ability to detect gene fusion without the decay as the coverage decline, indicating that 10× coverage was saturated for the two tools to identify fusions accurately (*Figure 2e-2f*). In contrast, JAFFAL and FusionSeeker exhibited notable precision degradation as coverage increased. For JAFFAL, precision declined from 79.9% to 75.0% for PacBio and from 78.9% to 75.3% for ONT. Similarly, FusionSeeker’s precision dropped from 68.3% to 62.0% for PacBio and from 68.5% to 62.2% for ONT. However, both tools maintained relatively stable sensitivities, with differences of at most 1.6% and 1.2% for JAFFAL, and 2.4% and 2.6% for FusionSeeker. We speculate that the lack of adaptive read support thresholds in JAFFAL and FusionSeeker leads to an increase in false positives as the number of reads grows. Remarkably, Genion showed a drastic decline in performance, with F1 scores dropping by 32.6% for PacBio and 23.6% for ONT when comparing 10× to 20× coverage data, highlighting Genion’s inability to effectively handle low-abundance transcripts, particularly at lower expressed transcripts.

We further evaluated these tools using negative simulations, which are based on a well-established background where few gene fusions are expected in non-tumor datasets. GFHunter reported the fewest fusions, with 7 (2 “Reliable” and 5 “Suspected”) and 6 (2 “Reliable” and 4 “Suspected”) fusions detected in the PacBio and ONT 30× datasets, respectively. Among these, 5 fusions were common between the two platforms, including 2 “Reliable” fusions (*Figure 2g, Supplementary Table 3*). Upon closer examination, the two “Reliable” fusions (GLCCI1-UMAD1 and EFCAB11-FOXN3) were found to be supported by lncRNA bridging the two genes, which is absent in current constructed reference transcriptome. By expanding the index from the protein-coding transcripts to include the full range of transcripts, GFHunter was able to avoid detecting any false positives in negative simulation datasets. LongGF and JAFFAL performed as the second and third best tools, identifying 10, 11 fusions for LongGF and 16, 15 fusions for JAFFAL in the PacBio and ONT datasets, respectively. In contrast, Genion and FusionSeeker produced significantly higher numbers of false positives, reporting at least 51 and 126 fusions, respectively. Notably, 76% of the fusions reported by FusionSeeker and Genion were supported by at least 8 reads, further highlighting their tendency to generate overabundant false positives. These results underscore GFHunter’s superior ability to effectively filter out false positives in negative simulated data, demonstrating its robustness.

### Fusion identification on real non-tumor data

Furthermore, we evaluated the ability of GFHunter to separate false positives on three real non-tumor datasets from a well-studied Genome in a Bottle human sample HG002. It was sequenced by PacBio Iso-seq, ONT dRNA and ONT cDNA technologies, with 4M, 12M, 57M throughput, 0.77%, 9.99%, 4.24% average error rate, and about 2539bp, 1286bp, 832bp N50 read length, respectively (*Supplementary Table 4*). The number of reported fusions is summarized in *Table 1*. For the two ONT datasets, GFHunter reported only 2 fusions each, both classified as “Reliable” with no overlapping fusions. JAFFAL reported one more high confidence fusion than GFHunter in the cDNA data, but 6 more in dRNA data. LongGF performed as the second-best tool, reporting only 5 false positives in the dRNA dataset. However, it identified an excessive number of fusions (42) in the cDNA dataset. A similar trend was observed with FusionSeeker, which reported 36 and 383 fusions in the dRNA and cDNA datasets, respectively. This over-reporting of fusions in cDNA data has been previously attributed to the creation of chimeric molecules during cDNA library preparation^39, 40^.

For PacBio Iso-seq data, GFHunter again demonstrated superior performance, reporting only 8 false positives classified as “Reliable”, followed by JAFFAL (16 false positives). Notably, PacBio data generally contained more potential fusions compared to ONT data (except FusionSeeker). We speculate that this is primarily due to the cDNA library preparation process also required for PacBio sequencing, which can introduce chimeric artifacts. Considering the lower rank fusions, GFHunter also reported the fewest false positives, but the increment was observed in cDNA and PacBio data, from 2 to 9, 8 to 16 for GFHunter and 3 to 24, 16 to 38 for JAFFAL. This trend further underscores the impact of chimeric molecule generation during cDNA library preparation and highlights the tendency of tools to call more false positives when using looser criteria with insufficient evidence. We further analyzed the overlap of fusions detected by the five tools across the three datasets. Remarkably, no fusion was reported by all tools simultaneously, and even when the threshold was reduced to 3 or 4 tools, only 2 fusions were pointed out. Additionally, the combinations of tools reporting these fusions varied significantly. Most fusions were detected independently by different tools, suggesting that genuine fusions are rare in non-tumor data and that the reported fusions are likely artifacts resulting from the sensitivity of each tool’s calling strategy or their tolerance for false signatures. Thus, GFHunter has an excellent ability to distinguish false positives and real gene fusions.

### Fusion identification on real cancer cell lines

While simulation studies provide valuable insights, they cannot fully capture the complexity and characteristics of real sequencing data from cancer samples. Factors such as the distribution and complexity of gene fusions, as well as the expression differences between fusion and normal transcripts, significantly impact the extraction of fusion signatures and the identification of fusion breakpoints. To further confirm GFHunter’s ability, six public long-read transcriptome sequencing datasets from three well-studied cancer cell lines, i.e., MCF-7, HCT-116 and SKBR-3 (*Supplementary Table 5*), where existing gene fusions had been proved though orthogonal validation of multiple computational approaches^35, 41^, or using RT-PCR^42^ and Sanger sequencing^41, 43^. MCF-7 is a human breast cancer cell line with estrogen, progesterone and glucocorticoid receptors^44^, datasets from PacBio Iso-seq, ONT cDNA and ONT dRNA sequencing technologies are analyzed. These datasets varied in total read coverage (1.8M, 6.8M, and 10.8M reads, respectively) and read identity (ranging from 14.66% to 29.87%) (*Figure 3a*). HCT-116 cell line was isolated from the colon of an adult male with colon cancer^45^, ONT cDNA and dRNA dataset were involved in the benchmark. The cDNA dataset was exceptionally high-expressed, containing 291M reads with 8.17% read identity. SKBR-3 cells were isolated from the pleural effusion of a female adenocarcinoma patient^46^, only a PacBio Iso-seq dataset is applied for the evaluation. Notably, the N50 read lengths of the PacBio Iso-seq datasets from MCF-7 (5,785 bp) and SKBR-3 (4,893 bp) were significantly longer than those of the ONT datasets, which were all below 1,500 bp (*Figure 3b*). The total number of fusions reported by each tool across the six datasets is summarized in *Supplementary Table 6*.

**Figure 3.**
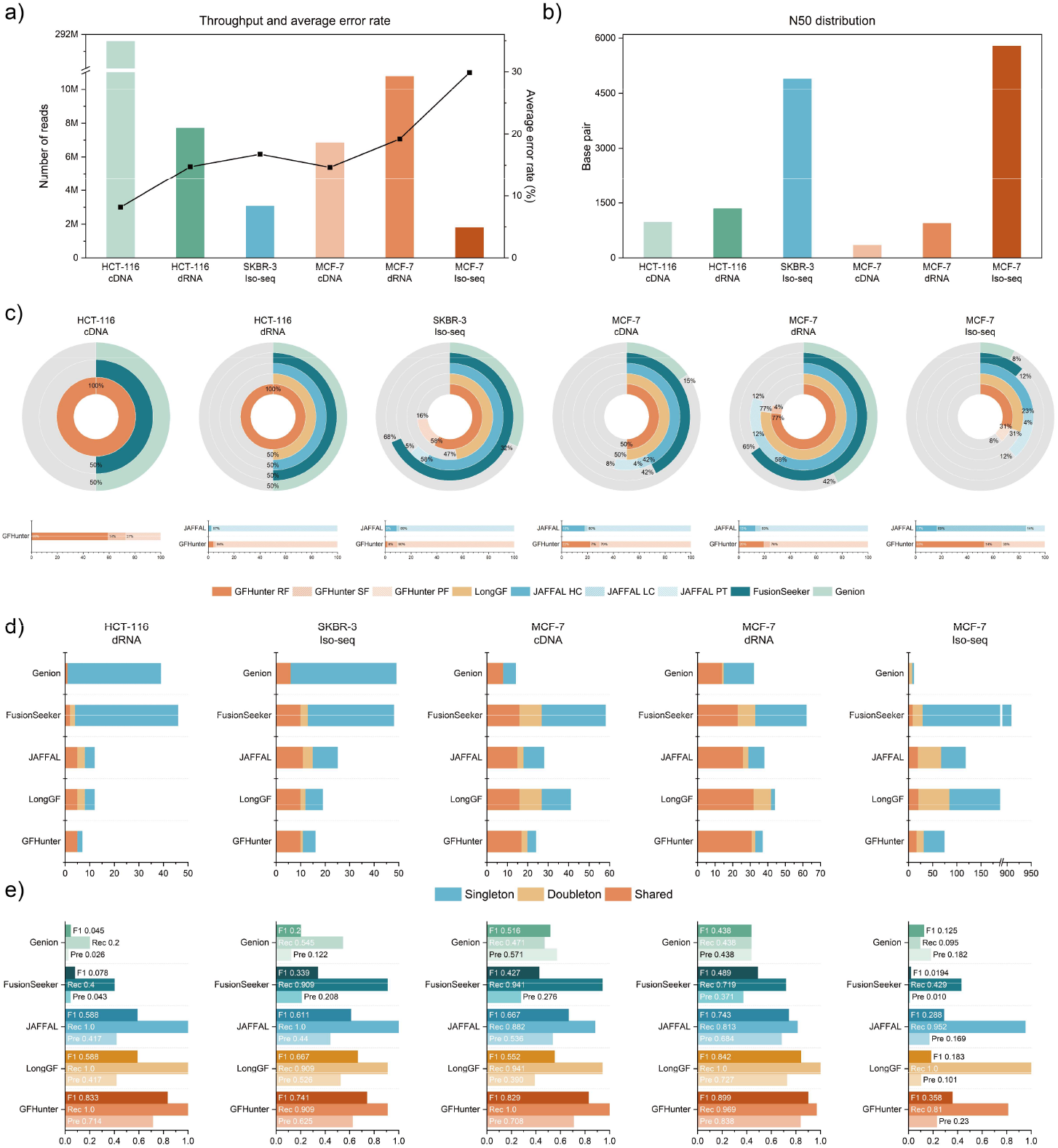
Benchmarking results on the real datasets. **a&b**. Throughput and sequencing error rates (a), and N50 length (b) for six real datasets (HCT-116 cDNA, HCT-116 dRNA, SKBR-3 Iso-seq, MCF-7 cDNA, MCF-7 dRNA, MCF-7 Iso-seq). **c**. The ratio of known fusion reported by different tools across various datasets (top panel), and the proportion of identified fusion with different ranks for GFHunter and JAFFAL (bottom panel). **d**. The number of Singleton, Doubleton and Shared fusions of each tool identified on different datasets. **e**. Recall (Rec), precision (Pre), and F1 score (F1) when using “Shared” fusions as the ground truth on the five datasets.

We first evaluated the ability to rediscover known fusions for different tools. Due to program crashes of LongGF and JAFFAL on the large HCT-116 cDNA dataset, their results were excluded. All the fusions with different ranks of GFHunter and JAFFAL are involved for the comparison. The discovery rates of known fusions across tools and datasets are shown in *Figure 3c* and *Supplementary Table 6-7*. GFHunter achieved the best discovery rates in most of the datasets (except MCF-7 cDNA dataset). Especially, GFHunter is only tool enable to report the two known fusions in HCT-116 cDNA and dRNA datasets, where SPAG9-MBTD1 fusion is uniquely called and classified as “Reliable”. The SPAG9-MBTD1 fusion^47^ associated with multiple cancers, was supported by only two split alignments (*Figure 4a* and *Supplementary Table 8*). In the IGV snapshot, one supporting read aligned to non-exon regions, likely due to sequencing errors or scoring system bias on alignment. However, such misalignments are avoided in GFHunter’s transcriptome-based approach, which focuses solely on transcript sequences, enabling the detection of this fusion. In SKBR-3 PacBio Iso-seq dataset, GFHunter uniquely identify another cancer-related “Potential” fusion CCDC85C-SETD3, known to influence cancer progression by altering cell proliferation and gene regulation pathways^42^. Despite clear alignment evidence from four supporting reads with homogeneous breakpoints in the IGV snapshot (*Figure 4b* and *Supplementary Table 8*), no other tools reported this fusion. Upon reviewing the alignments, we found that these four reads lacked split alignments because the fusion breakpoints were incorrectly connected and classified as intronic regions by the spliced alignment. This highlights the advantage of GFHunter’s transcriptome-based approach, which avoids such misclassifications. Although the fusion was categorized as 'Potential,' there is still a need for more expanded space to accurately recall key fusions. In absolute count, GFHunter and JAFFAL showed comparable fusion detection capabilities, with JAFFAL reporting one additional fusion (SMARCA4-CARM1) in the MCF-7 cDNA dataset. However, this fusion was supported by only a single read, suggesting potential over-sensitivity in JAFFAL’s calling strategy. FusionSeeker only report 3 fusions in MCF-7 PacBio Iso-seq data, likely due to the dataset’s high sequencing error rate (∼30%). Genion performed poorly across all datasets, indicating its limited suitability for long-read transcriptome fusion detection.

**Figure 4.**
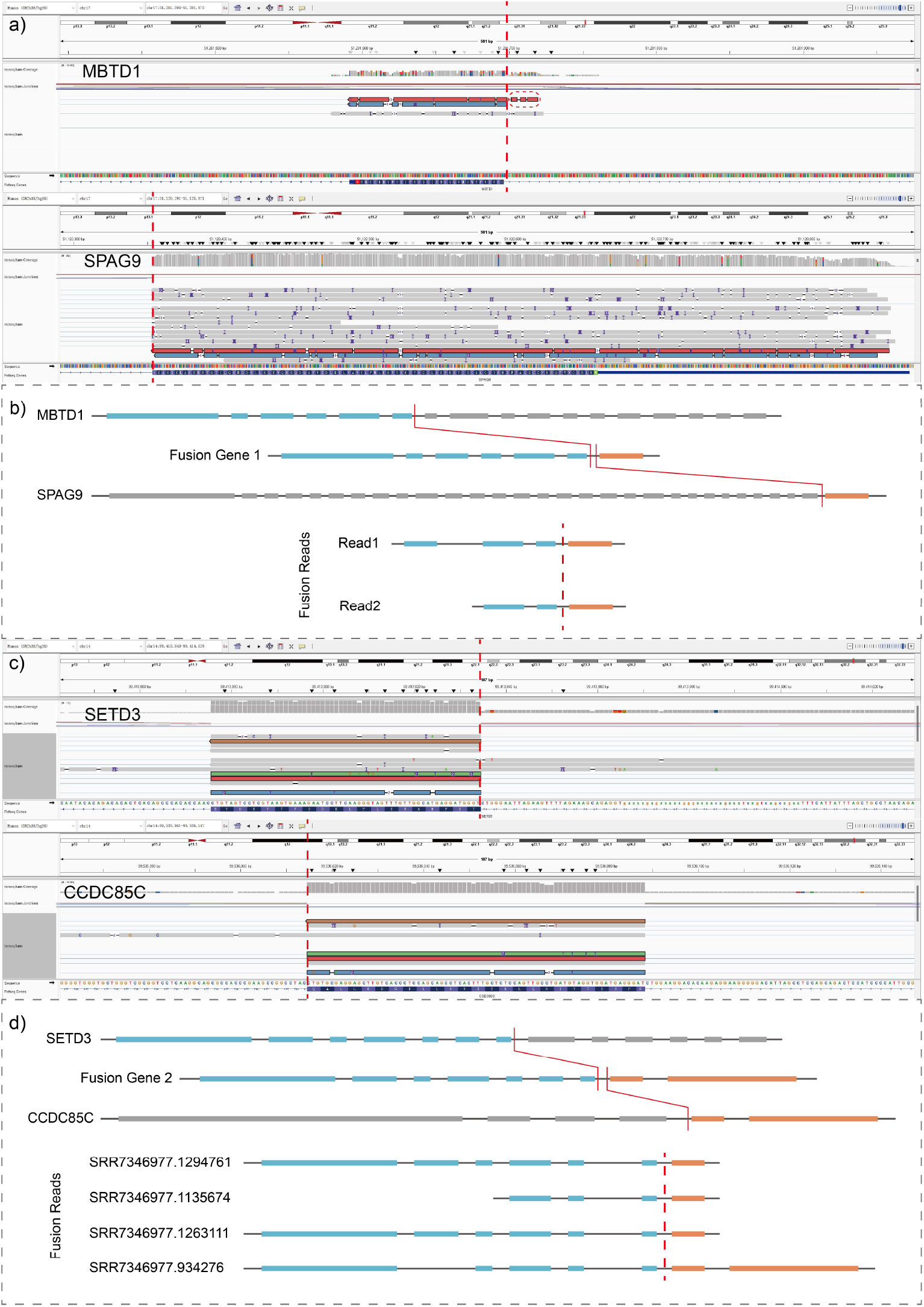
Schematic illustration of two known fusions of reported by GFHunter only. **a&b**. IGV snapshot of alignments (a) and illustration of exons in each read (b) supporting the fusion between genes MBTD1 (chr:17, 17 exons) and SPAG9 (chr:17, 30 exons). Only two reads are supported, with the red read aligning to a non-exon region (circled by the read frame). This results in the omission of this signaling read in genome-based alignment. **c&d**. IGV snapshot of alignments (c) and illustration of exons in each read (d) supporting the fusion between genes SETD3 (chr:14, 13 exons) and CCDC85C (chr:14, 6 exons). Four reads are aligned to the breakpoint near the exon boundary, but the inner part of the reads was misclassified as an intron by genome-based alignment.

While the above evaluation highlights the sensitivity of each tool in detecting known fusions, the assessment of false positives was not addressed. We observed that GFHunter’s “Reliable” fusions accounted for the majority of matched fusions (100%, 100%, 85%, 92%, 91%, and 79% across datasets). Similarly, JAFFAL’s “High Confidence” and “Low Confidence” fusions collectively showed comparable ratios (100%, 100%, 92%, 85%, 85%, and 69%). However, the total reported fusions across different confidence levels (bottom panel of *Figure 3c*) revealed that low-rank fusions constituted a significant proportion but rarely matched known fusions, suggesting a high rate of false positives. Therefore, we focused on GFHunter’s “Reliable” fusions and JAFFAL’s “High Confidence” and “Low Confidence” fusions for further precision and recall analysis.

Firstly, we categorized fusions into three groups: singleton (reported by 1 tool), doubleton (reported by 2 tools), and shared (reported by >2 tools). The distribution of these categories is shown in *Figure 3d* and *Supplementary Table 9* (excluding HCT-116 cDNA data). GFHunter achieved a higher proportion of shared fusions while simultaneously reporting fewer singleton and doubleton fusions across most datasets, demonstrating its superior capability to identify precise fusion events compared to competing tools. Specifically, GFHunter reported the most shared fusions (5 and 17) while exhibiting the fewest singleton fusions (2 and 4) in HCT-116 dRNA and MCF-7 cDNA datasets, respectively. Although LongGF detected slightly more shared fusions than GFHunter in the MCF-7 dRNA and Iso-seq datasets, it also reported significantly more doubleton and singleton fusions (6 for GFHunter vs. 12 for LongGF in MCF-7 dRNA, and 57 for GFHunter vs. 187 for LongGF in MCF-7 Iso-seq). Furthermore, Genion and FusionSeeker consistently produced a higher number of singleton fusions, suggesting a greater likelihood of false positives. Then, we conducted orthogonal validation using shared fusions as ground truth to evaluate the performance of each tool through precision, recall, and F1 scores. As illustrated in *Figure 3e* and *Supplementary Table 9*, GFHunter recorded the highest F1 scores across five datasets: HCT-116 dRNA (83.3%), SKBR-3 Iso-seq (74.1%), MCF-7 cDNA (82.9%), MCF-7 dRNA (89.9%), and MCF-7 Iso-seq (35.8%), surpassing the second-best tools by 24.5% (LongGF and JAFFAL), 7.4% (LongGF), 16.2% (JAFFAL), 5.7% (LongGF), and 7% (JAFFAL) respectively. GFHunter also demonstrated superior precision, highlighting its robust capability to accurately identify fusions from the shared callset. Notably, the F1 scores in the MCF-7 Iso-seq dataset were significantly lower due to poor accuracy, reflecting the challenges of limited shared and excessive singleton fusions. These results underscore GFHunter’s efficacy in accurately reporting tumor-associated known fusions, even with limited supportive evidence, outshining other state-of-the-art tools.

### Evaluation on the consistency of fusions across sequencing technologies

To further evaluate GFHunter’s ability to consistently detect and characterize fusions across different sequencing technologies, we performed a consistency analysis on three MCF-7 datasets: Iso-seq, cDNA, and dRNA (*Supplementary Table 10*). The upset plot in *Figure 5a* presents the counts and harmonic means (refer to *Online Method*) of consistently reported fusions compared to total reported fusions. Only the “Reliable” and “Suspected” fusions from GFHunter, and the “High Confidence” and “Low Confidence” fusions from JAFFAL, were considered, as false positives in the potential fusion calls were deemed irrelevant. Additionally, unvalidated fusions in the consistent set were excluded. Among fusions supported by both PacBio and ONT data, GFHunter (“Reliable” only) achieved the highest harmonic mean (13.3%, 6 fusions), indicating superior consistency compared to other tools. This was followed by GFHunter (“Reliable” + “Suspected”) at 10.8%, Genion at 10.5%, JAFFAL at 9.8%, LongGF at 6.1%, and FusionSeeker at 1.5%. Nevertheless, fusion consistency reported by tools across different sequencing protocols for the same sample was generally low. This may arise due to several factors: 1) discrepancies in error profiles and data quality across different sequencing technologies, 2) the inherent complexity of fusion genes, which include diverse fusion types, frequent alternative splicing, breakpoint variability, and differences in expression levels, 3) the varying effectiveness of different detection tools in identifying specific fusion events, and 4) batch effects present in the original sample preparations. When examining consistency for two shared fusions across the three datasets, GFHunter outperformed other tools, with nearly 50% harmonic means (49.2% for “Reliable”, 48.6% for “Reliable” + “Suspected”) between cDNA and dRNA datasets, demonstrating its ability to detect unified fusions. FusionSeeker (36.7%) and Genion (34.8%) exhibited the worst consistency. Notably, the high consistency observed in ONT data can be attributed to the similar sequencing principles and error patterns, with the main difference being the reverse transcription process in cDNA technology. Regarding consistency between PacBio and ONT technologies, GFHunter maintained superior performance, with harmonic means of 16.3% (between Iso-seq and cDNA, “Reliable”) and 14.4% (between Iso-seq and dRNA, “Reliable”), outperforming the other four tools despite the overall low absolute values.

**Figure 5.**
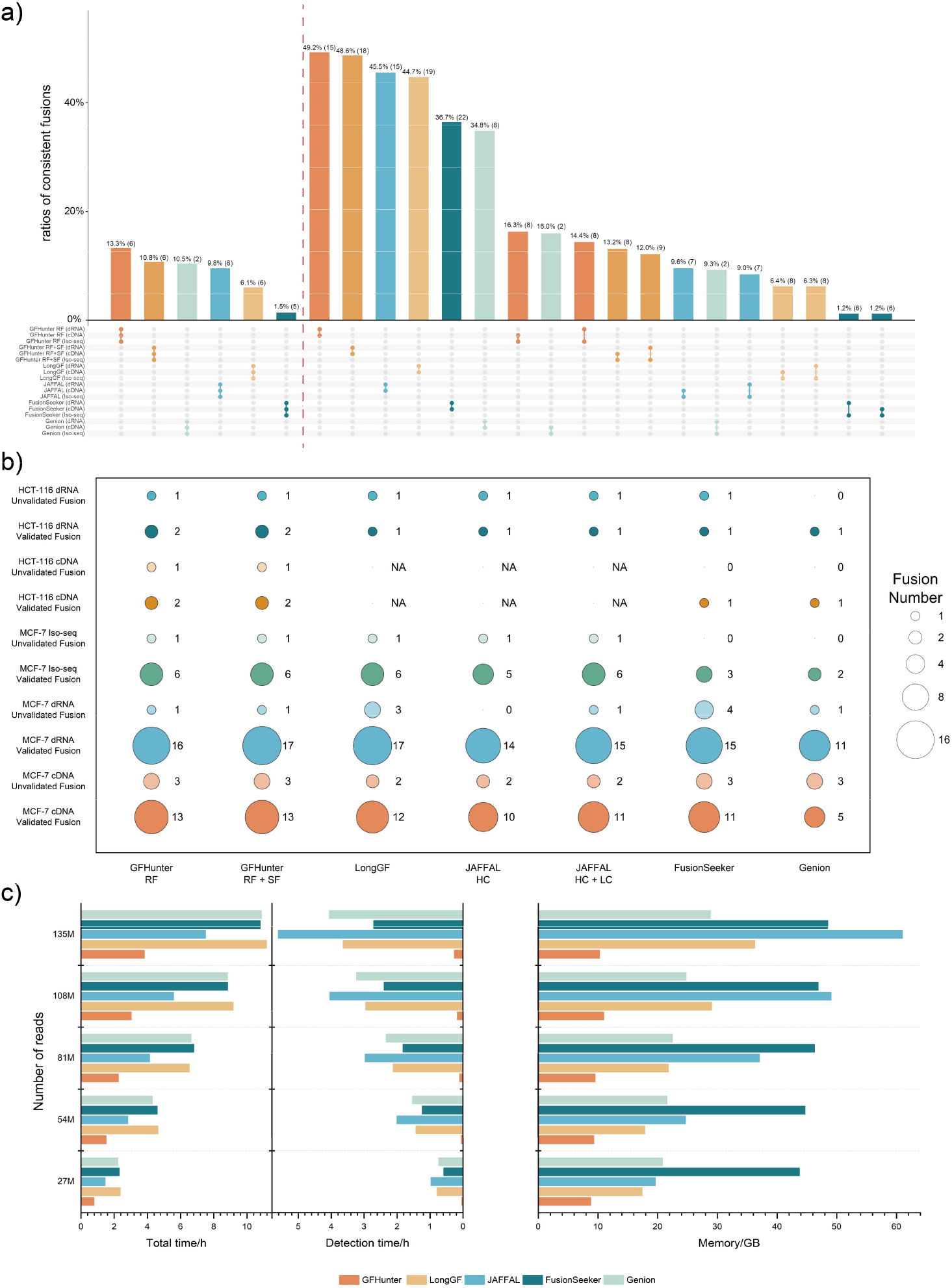
Benchmarking results on the consistency of fusions across sequencing technologies and Computational efficiency. **a**. Upset plot showing the harmonic mean and counts (in brackets) of consistently reported fusions of each tool on MCF-7 Iso-seq, dRNA and cDNA datasets. **b**. The number of consistent fusions reported by long-read and short-read sequencing data for the MCF-7 and HCT-116 cell lines across each tool. Fusions reported by short-read sequencing are treated as background and categorized into validated and unvalidated fusion groups. The results from long-read sequencing are then compared with these two groups, respectively (e.g., MCF-7 dRNA validated vs. MCF-7 dRNA unvalidated). GFHunter RF and SF represents the “Reliable” and “Suspected” fusion, respectively. And JAFFAL HC and LC represents the “High Confidence” and “Low Confidence” fusion, respectively. **c**. Runtime (left panel) and memory usage (right panel) under different input read counts for each tool. The runtime panel includes both the fusion detection time and the total time consumed.

Short-read transcriptome sequencing and various specialized tools have been widely used in cancer studies to identify gene fusions, which are potential markers for tumorigenesis and development^48^. We benchmarked GFHunter’s sensitivity by comparing fusion calls on matched long- and short-read data from the same samples, as genuine fusions should be detectable in both datasets. Two transcriptome datasets from MCF-7 and HCT-116 cancer cell lines, sequenced using Illumina technology, contained 152M and 137M reads, respectively. We used two popular short-read tools, Star-Fusion and JAFFA, to identify fusions, considering the jointly reported fusions as the background. In MCF-7 and HCT-116 short-read data, 24 and 4 fusions were detected, with 19 and 2 previously validated (*Supplementary Table 11*). In MCF-7 cDNA and dRNA data (*Figure 5b*), GFHunter reported 24 and 37 “Reliable” fusions, of which 16 and 17 were detected in the short-read data, achieving rediscovery rates of 66.7% and 45.9%, with 13 and 16 previously validated, respectively. LongGF and JAFFAL performed similarly to GFHunter, while Genion reported the fewest matching fusions. In PacBio data, although the number of detected fusions increased, the number of matches declined, indicating that PacBio’s sequencing technology is less effective than ONT in detecting fusions. In HCT-116 cDNA and dRNA data (*Figure 5b*), GFHunter matched 3 fusions, 2 of which were “Reliable”, while other tools matched at most 1 fusion. These results demonstrate GFHunter’s accuracy, stability across sequencing technologies, and the general utility of long-read data for fusion identification.

### Computational efficiency of GFHunter

We evaluated the speed and memory footprint of the fusion detection tools on a machine with 32 cores and 170 GB of memory. Five down-sampled datasets from the HCT-116 cDNA data, with varying total read coverages (27M, 54M, 81M, 108M, and 135M reads), were processed using 16 CPU cores. GFHunter was 2-3 times faster than other tools across all datasets when considering the entire workflow (alignment + fusion detection, referred to as *Total Time*) (*Figure 5c, Supplementary Table 12*). Focusing solely on fusion detection efficiency, GFHunter completed the task in less than 16 minutes for 135M reads, while other tools required at least 163 minutes (FusionSeeker). Overall, GFHunter was 10-30 times more efficient than other tools in parsing signatures and identifying fusions, primarily due to two key optimizations: 1) fast searching via an interval tree-based reference transcriptome index, and 2) efficient fusion gene pairing using sparse matrix multiplication. JAFFAL spent the most time on fusion detection but achieved the second-shortest total runtime, as it employs transcriptome-based alignment (similar to GFHunter), which is more efficient than genome-based alignment. However, JAFFAL’s runtime was significantly increased by the re-alignment of potential signal reads to the reference genome, a step that dominates its fusion calling process. Additionally, the runtime of all tools scaled linearly with the number of input reads.

Regarding memory usage, GFHunter consumed 1/6 to 1/2 of the memory required by other approaches, approximately 10GB, with only a slight increase in memory usage as the dataset size grew from 27M to 135M reads (*Figure 5c, Supplementary Table 12*). In contrast, FusionSeeker consistently required high memory usage, exceeding 40GB, while JAFFAL’s memory usage significantly increased from 19GB to 61GB, indicating its decreasing suitability for fusion calling on modern computing systems as read counts grow. We also evaluated the performance improvement with multi-threading on the SKBR-3 Iso-seq, using 1, 2, 4, and 8 CPU cores. GFHunter exhibited a near-linear speedup with the number of CPU cores, significantly reducing its wall clock time (*Supplementary Table 13*). In conclusion, GFHunter’s low resource requirements (both memory and time) make it a highly scalable fusion gene detection tool, well-suited for high-performance computing platforms and large-scale transcriptome data analysis tasks.

## Discussion

In this study, we explored the utility of long-read RNA sequencing for gene fusion detection, a critical aspect of oncology and genetic research. Long-read sequencing technologies, which generate significantly longer reads than traditional short-read platforms, offer substantial advantages in capturing full-length transcripts, enabling the detection of complex fusion events that may be missed by short-read sequencing due to its inherent limitations in read length.

However, it is not straightforward to detect gene fusions from noisy long-read transcriptome data, we propose GFHunter, a long-read fusion detection tool equipped with four specialized strategies for efficient and accurate fusion identification. First, GFHunter constructs and indexes a reference transcriptome using a hash-based approach, improving transcript diversity representation and minimizing alignment ambiguities. This enhances both accuracy and speed in the initial mapping, facilitating the identification of candidate fusions not present in a standard reference genome. Second, GFHunter identifies candidate gene pairs using sparse matrix multiplication, focusing on non-zero elements (potential fusion points) to optimize computational resources and scale efficiently across large datasets. Third, GFHunter clusters fusion breakpoints through bottom-up hierarchical clustering, followed by multiple sequence alignment to generate accurate fusion transcript sequences. This ensures a comprehensive understanding of fusion architecture, which is crucial for interpreting functional implications. Lastly, GFHunter refines and ranks each fusion event’s confidence level by cross-referencing results from both the transcriptome and genome, correcting misalignments in the initial mapping and ensuring precise breakpoint identification.

We benchmarked GFHunter on both simulated and real sequencing datasets, including non-tumor cell lines. Simulated data, mimicking ONT and PacBio platforms, demonstrated that GFHunter outperforms other tools in detecting spiked-in fusions, achieving the highest accuracy and F1 score. In real data from cancer cell lines, GFHunter reported the highest number of previously validated fusions, ranking them as “Reliable” with high confidence. Notably, GFHunter uniquely identified two fusions: SPAG9-MBTD1 in HCT-116 and CCDC85C-SETD3 in SKBR-3, critical in various cancers. These fusions, supported by only 2 and 4 split alignments, respectively, highlight GFHunter’s capability to rediscover fusions even with limited evidence. Orthogonal validation using shared fusions as ground truth further confirmed GFHunter’s superior performance, with F1 scores surpassing the second-best tools from 5.7% to 24.5%, emphasizing its sensitivity and accuracy. In datasets from non-tumor cell lines and negative simulations, where few fusions were expected, GFHunter consistently reported fewer “Reliable” fusions compared to other tools. This indicates a low false-positive rate, showcasing GFHunter’s excellent ability to distinguish between real gene fusions and artifacts.

Moreover, we found that cDNA data generally produce more fusions (potential false positives) than dRNA data, a trend more pronounced in PacBio data. This could be attributed to chimeric molecule formation during cDNA library preparation^39, 40^, leading to looser fusion detection criteria. Notably, GFHunter demonstrated the most consistent fusion calls across different sequencing protocols (long- and short-read), further underscoring the utility of long-read data for fusion identification. In the term of computational resource, GFHunter ran 2-3 times faster than other state-of-the-art tools across the entire calling workflow, and 10-30 times more efficient in parsing signatures and identifying fusions, using the least memory about 10GB that was 1/6 to 1/2 of other approaches, indicating the highly scalability of GFHunter.

Despite its strengths, GFHunter and similar alignment-and-signature-parsing methods have several limitations that warrant further improvement. 1) GFHunter filters out fusion transcripts without exon loss, excluding certain fusions, such as those caused by read-through transcription, to better differentiate between fusions resulting from DNA structural variations. 2) The consistency of fusion detection across different sequencing protocols remains suboptimal, even within ONT’s cDNA and dRNA technologies. This issue is not unique to GFHunter and highlights the challenges faced by all tools in ensuring consistent fusion detection. Notably, the differences in fusion calls across tools are well-documented, and an ensemble approach may be more effective for identifying actionable fusions, as it reduces tool- and data-specific biases. 3) GFHunter has yet to be optimized for single-cell long-read RNA-seq, which is crucial for accurately detecting fusions across diverse cell types. Although single-cell RNA-seq offers powerful insights into cell heterogeneity and transcript complexity, GFHunter’s current capabilities are limited. Future work will focus on developing a novel strategy for long-read single-cell fusion detection, distinct from approaches used in short-read sequencing^49^.

## Methods

GFHunter effectively identifies gene fusions in long-read transcriptome sequencing datasets through several specialized strategies, including reference transcriptome reconstruction and indexing, sparse matrix multiplication-based candidate gene pairing, bottom-up hierarchical clustering-based fusion breakpoints clustering, partial order alignment-based fusion transcript generation and cross-referencing-based fusion refinement and ranking. These methodologies consist of five key steps, detailed below (Figure 1).

### Reference transcriptome reconstruction and indexing

State-of-the-art approaches for long-read RNA-seq alignment typically map reads to either the reference genome or the reference transcriptome. The later exclusively preserves the sequence information within gene regions, that can expedite the alignment of spliced reads (also called “spliced alignment”), which typically consumes the majority of computational resources. However, reference transcriptome, such as those derived from GENCODE annotations, include a significant proportion of non-protein-coding transcripts—such as long non-coding RNAs, tRNAs, and transcripts from polymorphic pseudogenes. Lots of redundant or gene fusion unrelated transcripts may bring ambiguous alignments and lead to some false positive calls. To mitigate this issue, GFHunter reconstructs the reference transcriptome for efficient read alignment and accurate gene fusion detection.

The reconstruction of the reference transcriptome follows three steps:1) traversing the gene annotations and selectively preserving specific types of genes (primarily protein-coding genes preserved by default); 2) generating transcript sequences within each gene according to the splicing sites; and 3) concatenating the transcript sequences from all target genes on the same chromosome to generate a chromosome-level reference transcript. All the chromosome-level reference transcripts constitute the reconstructed reference transcriptome. To distinguish the transcripts that are concatenated, the offset and relevant annotations of each target transcript are recorded, generating the reconstructed annotations.

It is worth mentioning that, all the transcripts in the reconstructed reference transcriptome are in positive strand, which facilitates the identification of gene fusion signatures. Minimap2^50^ (optional) is used to index the reconstructed reference transcriptome for subsequent read alignment. Moreover, a hash-based index is employed to efficiently retrieve specific genes from any alignment position, primarily through the use of an interval tree. This enhances the speed and accuracy of gene identification during the alignment process.

### Transcriptome-based sequence alignment and filtering

Spliced aligners often struggle with short exons and numerous splicing events due to seed length limitations inherent in the mainstream “seeding-and-extension” alignment strategy. They also face challenges in identifying canonical and non-canonical splicing sites, especially with high sequencing errors or small variants. GFHunter overcomes these challenges by using transcriptome-based alignment instead of spliced alignment with Minimap2, where introns are excluded in the reconstructed reference transcriptome, simplifying the alignment process and improving accuracy. Additionally, the size of the reconstructed transcriptome is much smaller than that of the genome, reducing computational overhead.

GFHunter implements several advanced filters preserve potential gene fusion signatures for subsequent clustering analysis, including 1) excluding reads where the aligned portion is shorter than a specified minimum length (default: 50 bp); 2) retaining reads that map exclusively to a single gene and avoid large soft-clipping; 3) filtering reads where fusion breakpoints are located more than 30 bp away from exon-intron junctions. These stringent criteria effectively reduce the number of irrelevant reads, thereby enhancing the precision and efficiency of downstream gene fusion detection.

### Sparse matrix multiplication-based candidate gene pairing

The remaining reads, which align to at least two genes, require further analysis to determine the gene pairs involved in the fusion, based on their alignment patterns. This process is computationally intensive as it requires examining all possible gene pair combinations. To address this challenge, GFHunter utilizes sparse matrix multiplication to efficiently identify potential gene fusion pairs. This is achieved through Inner Product Optimization using NumPy, significantly enhancing the speed of candidate fusion pair detection.

Specifically, suppose there are *m* reads remaining and *n* genes identified though hash-based index. Each gene *g*_*i*_ is represented as an *m*-dimensional vector, where each element corresponds to a read. The value of an element is set to 1 if the read spans the gene, and 0 otherwise. This binary vector efficiently encodes gene-spanning information for each read as follow,

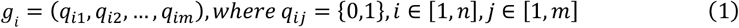

By arranging these *n* gene vectors, an *n* × *m* matrix *Q* is obtained.

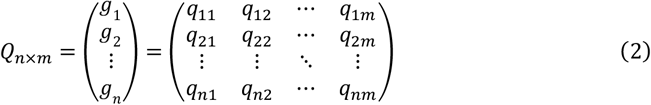

Since the matrix contains relatively few non-zero elements, it is considered a sparse matrix, allowing the use of sparse matrix storage and algorithms for efficient computation. Multiplying matrix *Q* with its transpose *Q*^*T*^ yields an *n* × *n* matrix *R*.

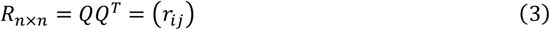

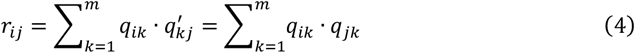

In matrix *R*, each off-diagonal element *r*_*ij*_ represents the number of reads supporting a fusion between gene *g*_*i*_ and *g*_*j*_. By traversing this symmetric matrix, gene pairs with at least two supporting reads (a user-defined parameter, default set to two) can be identified. For each identified gene pair, the corresponding vectors are compared element by element. If the values in a given position are the same (i.e., both 1), it indicates that the read supports the fusion between the two genes. This matrix *R* captures the co-occurrence patterns of reads spanning different gene pairs, which facilitate the identification of plausible gene fusions.

### Fusion scenarios recognition and breakpoint assignment

For each plausible gene fusion pair, four potential fusion types can arise, based on the relative positioning of genes *g*_1_ and *g*_2_ within the reconstructed reference transcriptome, split alignments, and the orientation of the alignment to the reference transcriptome. These possible fusion scenarios can be categorized, as summarized in *Supplementary Figure 1*. Therefore, GFHunter determines whether the breakpoint locate in front or behind of a specific exon, by evaluating the alignment flags and relative gene positions. Specifically, a breakpoint is considered to be behind a specific exon under the following conditions: (1) the gene is in the front, and the alignment flag equals 0, or (2) the gene is in the back, and the flag equals 16. Conversely, when (1) the gene is in the front, and the alignment flag equals 16, or (2) the gene is in the back, and the flag equals 0 is meet, the breakpoint is considered to be in front of a specific exon. Once the relative location of the breakpoint is determined, the distance between the fusion breakpoint and the corresponding exon boundary is calculated. If this distance exceeds a default threshold (30 bp), GFHunter discards the read as invalid. This ensures that only breakpoints located close enough to the exon boundaries are considered, enhancing the accuracy of fusion gene detection.

### Hierarchical clustering-based breakpoint clustering and fusion transcript generation

GFHunter organizes and standardizes each inferred fusion breakpoint pair as 2D coordinate points, which are then subjected to bottom-up hierarchical clustering based on Manhattan distances. Initially, each point is treated as an individual cluster. The hierarchical tree is constructed by iteratively merging the two closest clusters, determined by the distance between their centroids. The computation of Manhattan distances and cluster centroids follows the method described below:

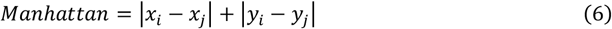

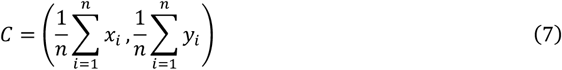

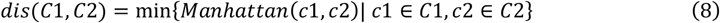

where (*x*_*i*_, *y*_*i*_) and (*x*_*j*_, *y*_*j*_) are the coordinates of two distinct fusion breakpoint pairs. A distance threshold *K* (default *K* = 200) is defined to identify hierarchical levels in the tree Specifically, when the distance between two clusters becomes less than *K*, they are considered to be merged (*Supplementary Figure 2*). Finally, clusters with at least 2 points are retained and others are discard. The coordinates are then de-standardized to restore the breakpoints on the reference genome. The bottom-up hierarchical clustering process allows for the identification of plausible fusion breakpoints within each cluster. Following this, the local sequences between the breakpoints are extracted from the corresponding reads. These extracted sequences are then aligned using multiple sequence alignment with abPOA to generate accurate fusion transcript sequence.

### Realignment and cross-referencing-based fusion event refinement and ranking

Transcriptome-based alignment depict the mapping signatures to the reconstructed reference transcriptome ignoring the search of splice junctions. But the ultimate goal is to identify the fusion breakpoints on the reference genome without bias. GFHunter calibrate and refine the fusion event and actual breakpoint by re-aligning the generated fusion transcript to the reference genome that considering the long-range alignment information with splicing site identification. GFHunter also ranks the confidence level of each fusion event by cross-referencing results from both the transcriptome and genome. Three main scenarios are considered:

**Scenario 1**. If the fusion transcript cannot be aligned to the reference genome or lacks a reliable supplementary alignment, the fusion event is deemed invalid and classified as a “Potential” fusion. Such candidates are subsequently excluded from further analysis.

This scenario may occur due to two primary reasons. Firstly, the presence of soft-clipped regions in the reads, which are too short to provide sufficient evidence for split alignment, can hinder the accurate identification of fusion breakpoints. Secondly, the inherent differences between genome and transcriptome representations: while the genome sequence reflects the linear order of exons within genes, the transcriptome reveals the combinatorial arrangement of exons. This allows transcriptome-based alignment to bypass the need for splicing site and exon recognition, potentially leading to discrepancies.

**Scenario 2**. If a supplementary alignment is present, but the gene it maps to differs from the original fusion gene pair, or if the supplementary alignment maps to the same gene but the breakpoint difference exceeds a predefined threshold *k* (default *k* = 200)., the fusion event is classified as a “Suspected” fusion, indicating a potential false positive.

**Scenario 3**. If the fusion event and breakpoint identified through realignment are identical to those derived from transcriptome-based alignment, the fusion gene pair is confirmed as a “Reliable” fusion.

### Metrics for the evaluation

The precision, recall and F1 score are used to evaluate the reported fusions comparing to the ground truth, which are defined as follows:

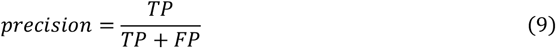

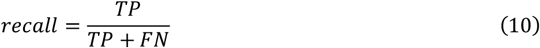

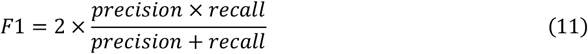

where TP (true positive) and FP (false positive) represent the number of fusions that match/mismatch the ground truth. FN (false negative) represent the number of fusions in the ground truth that are unreported.

The consistency of reported fusions between different sequencing datasets for a specific tool is evaluated using harmonic mean using the ratios of consistent fusions divide the number of totally report fusion, as follow described:

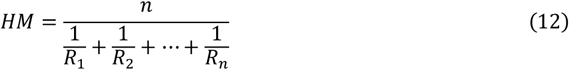

In this context, *n* is the number of datasets, and *R*_*i*_ is the ratio of *i*-th dataset.

All the benchmarks were implemented using a server with 2 Intel(R) Xeon(R) Gold 6240 CPUs @ 2.60 GHz (32 cores in total), 128 gigabytes of RAM, running on the CentOS Linux release 7.5.1804 operating system. The elapsed time and memory footprint were assessed by using the “seff” command of the Slurm Workload Manager.

## Data availability

The links to the reference genome, gene annotation, public datasets used for benchmarking are available in *Supplementary Table 14*. All the version and download URL of tools used in this study are listed in *Supplementary Table 15*. All the reported fusions in this study are publicly available at https://github.com/luzhenhao-HIT/GFHunter. The simulation workflow, the benchmarked commands, the fusion detection commands and related benchmarking materials used in this study are available in *Supplementary Notes*.

## Code availability

The GFHunter was implemented in Python and can be easily installed via PyPI. Its source code is available at https://github.com/luzhenhao-HIT/GFHunter. The codes support the interface to generate user’s specific reference transcriptome with a given annotation.

## Declarations

### Consent for publication

Not applicable

### Author contributions

YL and ZL designed the method. ZL implemented the method and performed the experiments and data analysis. YL and ZL wrote the manuscript. YL and ZL contributed equally to this work. The authors read and approved the final manuscript.

