## Supplementary Material for "GFHunter enables accurate and efficient gene fusion detection in long-read cancer transcriptomes"

### Supplementary Materials

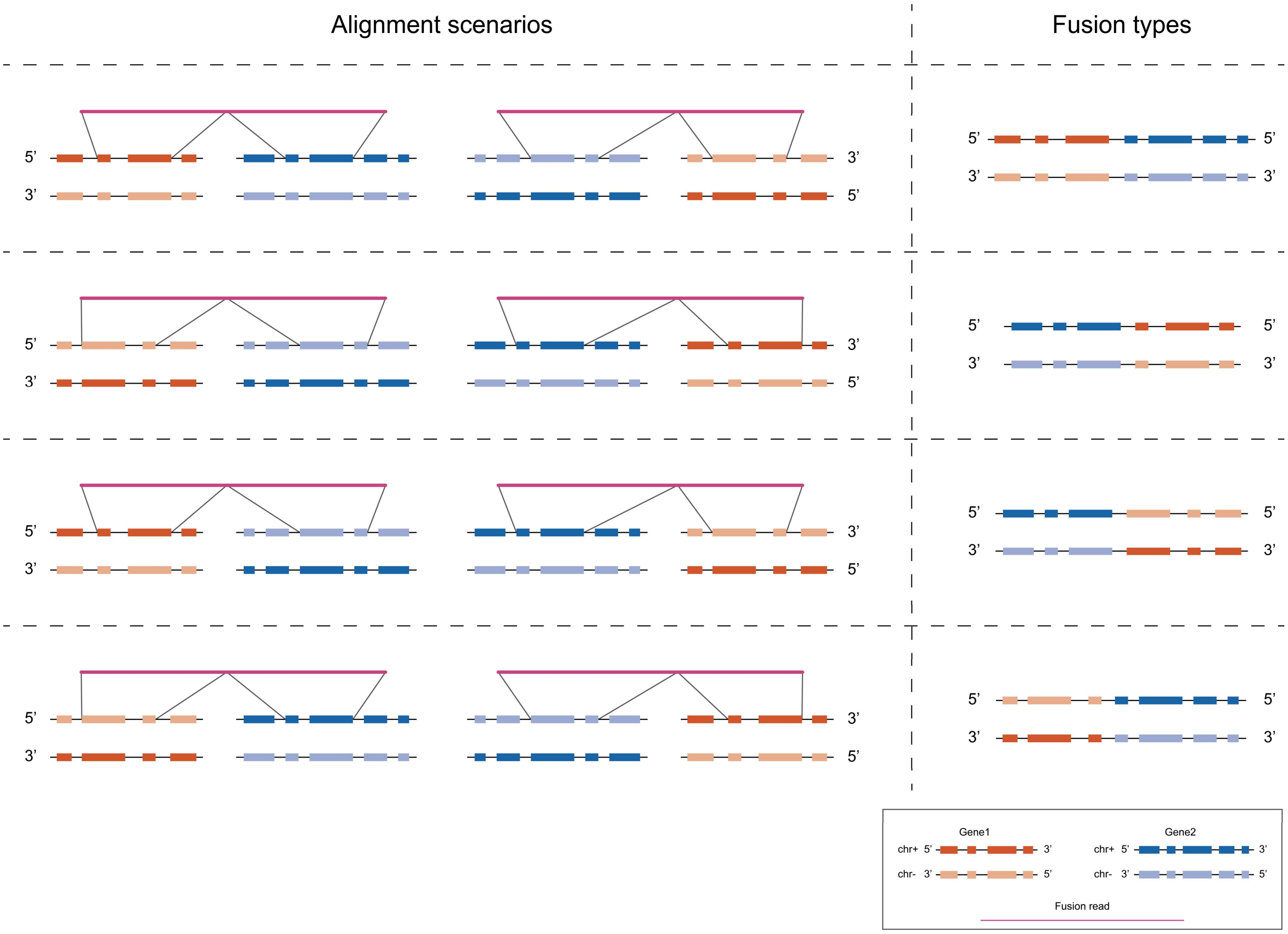

**Supplementary Figure 1. Four potential fusion types and corresponding alignment scenarios.**

Each fusion type encompasses two scenarios: one derived from a positive-strand fusion transcript and the other from a negative-strand fusion transcript. GFHunter determines the specific fusion type by analyzing alignment results, such as the 2×4 scenarios, to accurately calculate the true breakpoint positions in the genome. This approach ensures precise identification and localization of fusion events.

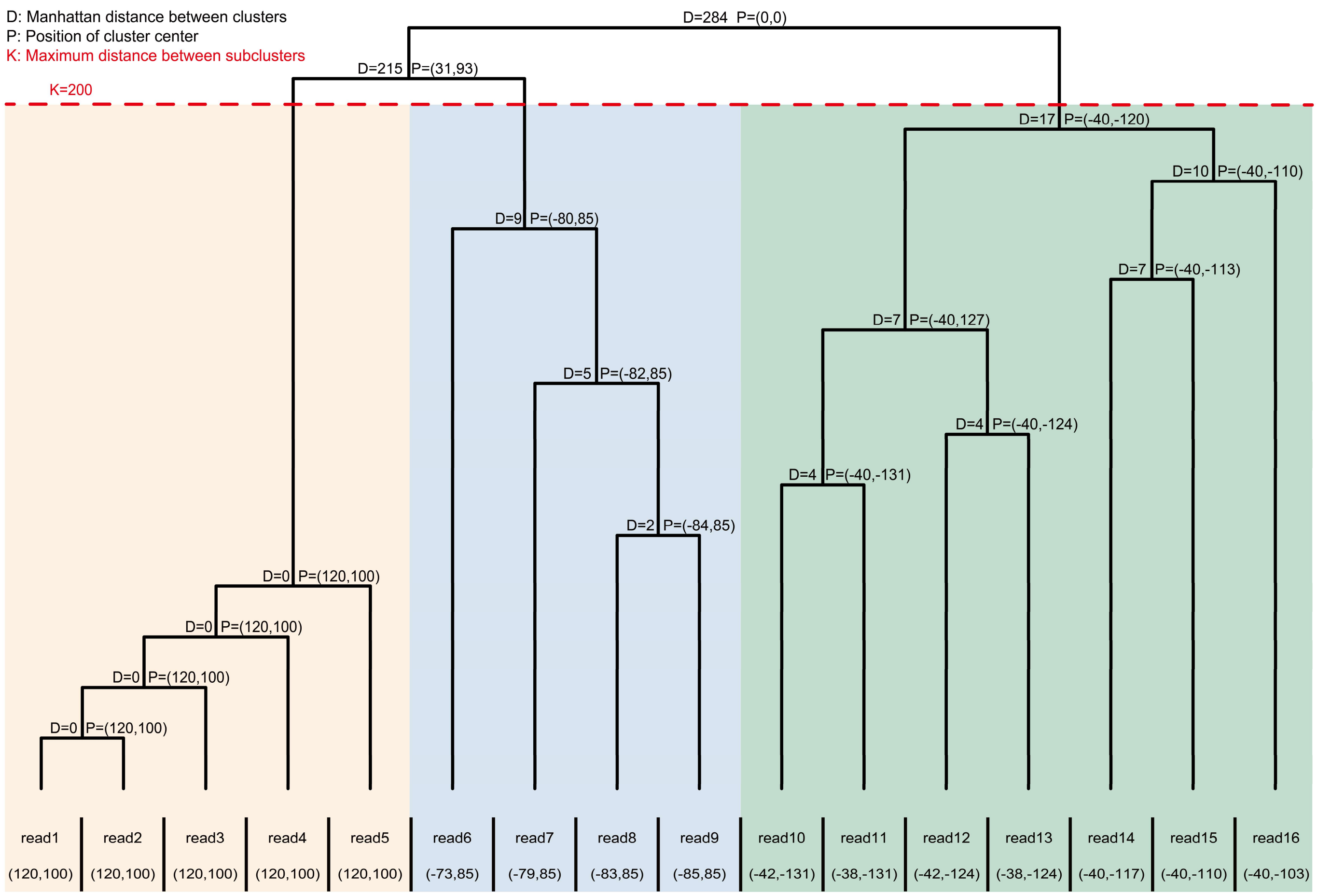

**Supplementary Figure 2. An example of bottom-up clustering.**

An example of bottom-up clustering is demonstrated with 16 reads that have standardized coordinates. Initially, each read is treated as an individual subcluster. In the clustering process, the two nearest subclusters are merged into a new cluster, repeating this process until the distance between the closest subclusters exceeds a predefined threshold $K$. In this example, with $K = 200$, the reads are ultimately grouped into three clusters: read1 to read5 form the first cluster, read6 to read9 form the second cluster, and read10 to read16 form the third cluster.

**Supplementary Table 1. The source of positive simulations.**

| chr | 1 | 2 | 3 | 4 | 5 | 6 | 7 | 8 | 9 | 10 | 11 | 12 | 13 | 14 | 15 | 16 | 17 | 18 | 19 | 20 | 21 | 22 | X |
|---|---|---|---|---|---|---|---|---|---|---|---|---|---|---|---|---|---|---|---|---|---|---|---|
| **1** | 5 | 5 | 5 | 5 | 4 | 3 | 6 | 4 | 5 | 7 | 10 | 3 | 1 | 1 | 1 | 6 | 6 | 1 | 3 | 1 | 3 | 1 | 3 |
| **2** | | 2 | 8 | 3 | 3 | 3 | 2 | 1 | 1 | 1 | 5 | 5 | 2 | 5 | 3 | 5 | 5 | 1 | 5 | 1 | 0 | 0 | 1 |
| **3** | | | 1 | 1 | 1 | 5 | 4 | 2 | 2 | 1 | 1 | 2 | 0 | 1 | 3 | 4 | 1 | 1 | 5 | 1 | 0 | 3 | 4 |
| **4** | | | | 1 | 0 | 3 | 2 | 3 | 3 | 1 | 3 | 0 | 0 | 0 | 0 | 1 | 5 | 1 | 3 | 2 | 0 | 0 | 1 |
| **5** | | | | | 1 | 2 | 2 | 1 | 2 | 0 | 5 | 2 | 1 | 2 | 1 | 2 | 5 | 2 | 6 | 0 | 0 | 2 | 3 |
| **6** | | | | | | 3 | 4 | 1 | 0 | 2 | 2 | 1 | 0 | 1 | 1 | 2 | 2 | 0 | 2 | 2 | 0 | 2 | 0 |
| **7** | | | | | | | 2 | 2 | 3 | 2 | 1 | 5 | 1 | 2 | 0 | 7 | 1 | 1 | 5 | 1 | 0 | 1 | 0 |
| **8** | | | | | | | | 2 | 2 | 3 | 5 | 1 | 1 | 2 | 5 | 1 | 3 | 1 | 4 | 1 | 0 | 1 | 1 |
| **9** | | | | | | | | | 3 | 1 | 1 | 1 | 2 | 2 | 4 | 0 | 2 | 0 | 3 | 1 | 0 | 0 | 1 |
| **10** | | | | | | | | | | 1 | 0 | 1 | 0 | 2 | 0 | 0 | 1 | 0 | 0 | 0 | 0 | 1 | 2 |
| **11** | | | | | | | | | | | 1 | 6 | 0 | 1 | 1 | 2 | 7 | 1 | 3 | 2 | 2 | 0 | 1 |
| **12** | | | | | | | | | | | | 0 | 0 | 2 | 3 | 1 | 3 | 0 | 2 | 0 | 2 | 0 | 1 |
| **13** | | | | | | | | | | | | | 0 | 1 | 1 | 0 | 2 | 1 | 0 | 0 | 0 | 0 | 0 |
| **14** | | | | | | | | | | | | | | 1 | 2 | 1 | 1 | 0 | 6 | 0 | 0 | 2 | 1 |
| **15** | | | | | | | | | | | | | | | 1 | 3 | 0 | 2 | 3 | 0 | 0 | 4 | 3 |
| **16** | | | | | | | | | | | | | | | | 0 | 4 | 2 | 3 | 3 | 0 | 0 | 3 |
| **17** | | | | | | | | | | | | | | | | | 1 | 1 | 3 | 1 | 0 | 2 | 4 |
| **18** | | | | | | | | | | | | | | | | | | 0 | 0 | 1 | 0 | 0 | 0 |
| **19** | | | | | | | | | | | | | | | | | | | 3 | 0 | 1 | 1 | 4 |
| **20** | | | | | | | | | | | | | | | | | | | | 0 | 1 | 0 | 0 |
| **21** | | | | | | | | | | | | | | | | | | | | | 0 | 0 | 0 |
| **22** | | | | | | | | | | | | | | | | | | | | | | 0 | 1 |
| **X** | | | | | | | | | | | | | | | | | | | | | | | 2 |

**Supplementary Table 2. The benchmarking result on simulated datasets.**

| | TP | FP | FN | Recall | Precision | F1-score |
|---|---|---|---|---|---|---|
| **Simulation PacBio 10x** | | | | | | |
| GFHunter[a] | 469 | 14 | 31 | 0.938 | 0.971 | 0.954 |
| LongGF | 475 | 21 | 25 | 0.950 | 0.958 | 0.954 |
| JAFFAL[b] | 393 | 99 | 107 | 0.786 | 0.799 | 0.792 |
| FusionSeeker | 399 | 185 | 101 | 0.798 | 0.683 | 0.736 |
| Genion | 155 | 50 | 345 | 0.310 | 0.756 | 0.440 |
| **Simulation PacBio 20x** | | | | | | |
| GFHunter[a] | 474 | 12 | 26 | 0.948 | 0.975 | 0.961 |
| LongGF | 489 | 27 | 11 | 0.978 | 0.948 | 0.963 |
| JAFFAL[b] | 397 | 109 | 103 | 0.794 | 0.785 | 0.789 |
| FusionSeeker | 407 | 219 | 93 | 0.814 | 0.650 | 0.723 |
| Genion | 359 | 78 | 141 | 0.718 | 0.822 | 0.766 |
| **Simulation PacBio 30x** | | | | | | |
| GFHunter[a] | 478 | 17 | 22 | 0.956 | 0.966 | 0.961 |
| LongGF | 489 | 33 | 11 | 0.978 | 0.937 | 0.957 |
| JAFFAL[b] | 396 | 117 | 104 | 0.792 | 0.772 | 0.782 |
| FusionSeeker | 410 | 233 | 90 | 0.820 | 0.638 | 0.717 |
| Genion | 405 | 91 | 95 | 0.810 | 0.817 | 0.813 |
| **Simulation PacBio 50x** | | | | | | |
| GFHunter[a] | 482 | 16 | 18 | 0.964 | 0.968 | 0.966 |
| LongGF | 491 | 38 | 9 | 0.982 | 0.928 | 0.954 |
| JAFFAL[b] | 401 | 134 | 99 | 0.802 | 0.750 | 0.775 |
| FusionSeeker | 411 | 252 | 89 | 0.822 | 0.620 | 0.707 |
| Genion | 415 | 104 | 85 | 0.830 | 0.800 | 0.815 |
| **Simulation ONT 10x** | | | | | | |
| GFHunter[a] | 481 | 19 | 19 | 0.962 | 0.962 | 0.962 |
| LongGF | 489 | 28 | 11 | 0.978 | 0.946 | 0.962 |
| JAFFAL[b] | 399 | 107 | 101 | 0.798 | 0.789 | 0.793 |
| FusionSeeker | 398 | 183 | 102 | 0.796 | 0.685 | 0.736 |
| Genion | 222 | 56 | 278 | 0.444 | 0.799 | 0.571 |
| **Simulation ONT 20x** | | | | | | |
| GFHunter[a] | 483 | 20 | 17 | 0.966 | 0.960 | 0.963 |
| LongGF | 492 | 25 | 8 | 0.984 | 0.952 | 0.968 |
| JAFFAL[b] | 398 | 115 | 102 | 0.796 | 0.776 | 0.786 |
| FusionSeeker | 408 | 211 | 92 | 0.816 | 0.659 | 0.729 |
| Genion | 388 | 74 | 112 | 0.776 | 0.840 | 0.807 |
| **Simulation ONT 30x** | | | | | | |
| GFHunter[a] | 486 | 19 | 14 | 0.972 | 0.962 | 0.967 |
| LongGF | 489 | 36 | 11 | 0.978 | 0.931 | 0.954 |
| JAFFAL[b] | 400 | 117 | 100 | 0.800 | 0.774 | 0.787 |
| FusionSeeker | 410 | 224 | 90 | 0.820 | 0.647 | 0.723 |
| Genion | 413 | 89 | 87 | 0.826 | 0.823 | 0.824 |
| **Simulation ONT 50x** | | | | | | |
| GFHunter[a] | 484 | 17 | 16 | 0.968 | 0.966 | 0.967 |

| | TP | FP | FN | Recall | Precision | F1-score |
|---|---|---|---|---|---|---|
| LongGF | 490 | 36 | 10 | 0.980 | 0.932 | 0.955 |
| JAFFAL[b] | 405 | 133 | 95 | 0.810 | 0.753 | 0.780 |
| FusionSeeker | 411 | 250 | 89 | 0.822 | 0.622 | 0.708 |
| Genion | 420 | 102 | 80 | 0.840 | 0.805 | 0.822 |

a: “Reliable” and “Suspected” fusions are used for GFHunter;

b: “High Confidence” and “Low Confidence” fusions are used for JAFFAL.

**Supplementary Table 3. The reported fusions on negative simulated datasets.**

| Datasets | GFHunter RF | GFHunter SF | LongGF | JAFFAL HC | JAFFAL LC | FusionSeeker | Genion |
|---|---|---|---|---|---|---|---|
| PacBio 30× | 2 | 5 | 10 | 10 | 6 | 126 | 51 |
| ONT 30× | 2 | 4 | 11 | 11 | 4 | 115 | 50 |

RF: “Reliable” fusions; SF: “Suspected” fusions; HC: “High Confidence”; LC: “Low Confidence”.

**Supplementary Table 4. Description of real non-tumor data (HG002).**

| Datasets | Throughput (#read) | Average error rate | N50 read length (bp) |
|---|---|---|---|
| PacBio Iso-seq | 4,434,605 | 0.77% | 2,539 |
| ONT dRNA | 12,067,989 | 9.99% | 1,286 |
| ONT cDNA | 57,355,400 | 4.24% | 832 |

**Supplementary Table 5. Description of real cancer cell lines.**

| Dataset | Throughput (#read) | Average error rate | N50 read length (bp) |
|---|---|---|---|
| HCT-116 cDNA | 291,592,884 | 8.17% | 973 |
| HCT-116 dRNA | 7,712,110 | 14.74% | 1,343 |
| SKBR-3 Iso-seq | 3,070,545 | 16.76% | 4,893 |
| MCF-7 cDNA | 6,837,532 | 14.66% | 714 |
| MCF-7 dRNA | 10,759,489 | 19.21% | 942 |
| MCF-7 Iso-seq | 1,789,323 | 29.87% | 5,785 |

**Supplementary Table 6. The total number of fusions reported by each tool.**

| | GFHunter | | | LongGF | JAFFAL | | | FusionSeeker | Genion |
|---|---|---|---|---|---|---|---|---|---|
| | RF | SF | PF | | HC | LC | PT | | |
| HCT-116 cDNA | 3359 | 769 | 1530 | NA | NA | NA | NA | 538 | 424 |
| HCT-116 dRNA | 7 | 3 | 156 | 12 | 9 | 3 | 388 | 46 | 38 |
| SKBR-3 Iso-seq | 16 | 67 | 789 | 19 | 20 | 5 | 199 | 48 | 48 |
| MCF-7 cDNA | 24 | 8 | 76 | 41 | 25 | 3 | 111 | 58 | 13 |
| MCF-7 dRNA | 37 | 9 | 143 | 44 | 29 | 9 | 183 | 62 | 31 |
| MCF-7 Iso-seq | 74 | 19 | 46 | 208 | 23 | 95 | 20 | 908 | 10 |

RF: “Reliable” fusions; SF: “Suspected” fusions; PF: “Potential” fusions;

HC: “High Confidence”; LC: “Low Confidence”; PT: “Potential Trans-Splicing”

**Supplementary Table 7. The number of previously validated fusions rediscovered across six long-read sequencing datasets.**

| Dataset | Fusion genes validated | GFHunter | | | LongGF | JAFFAL | | | FusionSeeker | Genion |
|---|---|---|---|---|---|---|---|---|---|---|
| | | RF | SF | PF | | HC | LC | PT | | |
| HCT-116 cDNA | COMMD10:AP3S1 | √ | | | | | | | √ | √ |
| | SPAG9:MBTD1 | √ | | | | | | | | |
| | Total | 2 | 0 | 0 | NA | NA | NA | NA | 1 | 1 |
| HCT-116 dRNA | COMMD10:AP3S1 | √ | | | √ | √ | | | √ | √ |
| | SPAG9:MBTD1 | √ | | | | | | | | |
| | Total | 2 | 0 | 0 | 1 | 1 | 0 | 0 | 1 | 1 |
| SKBR-3 Iso-seq | TATDN1:GSDMB | | | √ | √ | √ | | | √ | |
| | RARA:PKIA | √ | | | | | | √ | | |
| | CCDC85C:SETD3 | | | √ | | | | | | |
| | SUMF1:LRRFIP2 | √ | | | √ | √ | | | √ | √ |
| | TBC1D31:ZNF704 | √ | | | √ | √ | | | √ | |
| | CYTH1:EIF3H | √ | | | | √ | | | | |
| | DHX35:ITCH | √ | | | √ | √ | | | √ | √ |
| | NFS1:PREX1 | | | | | | | | | |
| | ATAD5:TLK2 | √ | | | √ | √ | | | √ | √ |
| | COL14A1:MTSS1 | | | | | | | | | |
| | DEPDC1B:PDE4D | √ | | | √ | √ | | | √ | |
| | PBRM1:WDR82 | √ | | | √ | √ | | | | |
| | PREX1:CPNE1 | | | √ | | | | | √ | |
| | TAF2:COLEC10 | √ | | | √ | √ | | | √ | √ |
| | TRIO:FBXL7 | √ | | | | √ | | | √ | √ |
| | KLHDC2:SNTB1 | √ | | | √ | √ | | | √ | √ |
| | SAMD12:MTBP | | | | | | | | √ | |
| | RAD51B:SEMA6D | | | | | | | | √ | |
| | TOX2:STAU1 | | | | | | | | √ | |
| | Total | 11 | 0 | 3 | 9 | 11 | 0 | 1 | 13 | 6 |
| MCF-7 cDNA | BCAS4:BCAS3 | √ | | | √ | √ | | | √ | √ |
| | ARFGEF2:SULF2 | √ | | | √ | √ | | | √ | |
| | RPS6KB1:VMP1 | √ | | | √ | √ | | | √ | |
| | SMARCA4:CARM1 | | | | | | | | | |
| | SLC25A24:NBPF6 | √ | | | √ | | √ | | √ | √ |
| | TBL1XR1:RGS17 | | | | √ | √ | | | | |
| | RPS6KB1:DIAPH3 | | | | | | | | | |
| | AHCYL1:RAD51C | √ | | | √ | √ | | | √ | |
| | TXLNG:SYAP1 | √ | | | √ | √ | | | √ | √ |
| | MYO6:SENP6 | √ | | | √ | √ | | | √ | |
| | POP1:MATN2 | √ | | | | | | √ | | |
| | GATAD2B:NUP210L | √ | | | √ | √ | | | √ | |
| | ESR1:CCDC170 | | | | | | | | | |
| | DEPDC1B:ELOVL7 | √ | | | √ | √ | | | √ | |
| | ATXN7L3:FAM171A2 | | | | | | | | | |
| | SYTL2:PICALM | | | | | | | | | |

| Dataset | Fusion genes validated | GFHunter | | | LongGF | JAFFAL | | | FusionSeeker | Genion |
|---|---|---|---|---|---|---|---|---|---|---|
| | | RF | SF | PF | | HC | LC | PT | | |
| | MYO9B:FCHO1 | | | | | | | | | |
| | PAPOLA:AK7 | √ | | | √ | | | √ | | |
| | ATP1A1:ZFP64 | √ | | | √ | √ | | | √ | |
| | BCAS3:ATXN7 | | | | | | | | | |
| | BCAS4:ZMYND8 | √ | | | √ | √ | | | √ | √ |
| | NAV1:GPR37L1 | | | | | | | | | |
| | NCOA3:SULF2 | | | | | | | | | |
| | PLCG1:TOP1 | | | | | | | | | |
| | PNPLA7:DPH7 | | | | | | | | | |
| | SULF2:PRICKLE2 | | | | | | | | | |
| | Total | 13 | 0 | 0 | 13 | 11 | 1 | 2 | 11 | 4 |
| MCF-7 dRNA | BCAS4:BCAS3 | √ | | | √ | √ | | | √ | √ |
| | ARFGEF2:SULF2 | √ | | | √ | √ | | | √ | √ |
| | RPS6KB1:VMP1 | √ | | | √ | √ | | | √ | √ |
| | SMARCA4:CARM1 | √ | | | √ | √ | | | √ | |
| | SLC25A24:NBPF6 | √ | | | √ | √ | | | √ | √ |
| | TBL1XR1:RGS17 | √ | | | √ | √ | | | √ | √ |
| | RPS6KB1:DIAPH3 | | | | | | | | | |
| | AHCYL1:RAD51C | √ | | | √ | √ | | | √ | |
| | TXLNG:SYAP1 | √ | | | √ | √ | | | √ | √ |
| | MYO6:SENP6 | √ | | | √ | √ | | | √ | |
| | POP1:MATN2 | | | | | | | | | |
| | GATAD2B:NUP210L | √ | | | √ | √ | | | √ | √ |
| | ESR1:CCDC170 | | | | | | | | | |
| | DEPDC1B:ELOVL7 | √ | | | √ | √ | | | √ | √ |
| | ATXN7L3:FAM171A2 | √ | | | √ | √ | | | √ | √ |
| | SYTL2:PICALM | √ | | | √ | √ | | | √ | √ |
| | MYO9B:FCHO1 | √ | | | √ | √ | | | √ | |
| | PAPOLA:AK7 | | √ | | √ | | | √ | √ | |
| | ATP1A1:ZFP64 | √ | | | √ | √ | | | √ | |
| | BCAS3:ATXN7 | | | | | | | √ | √ | |
| | BCAS4:ZMYND8 | √ | | | √ | | √ | | | |
| | NAV1:GPR37L1 | √ | | | √ | | | √ | | √ |
| | NCOA3:SULF2 | | | | | | | | | |
| | PLCG1:TOP1 | √ | | | √ | | √ | | | |
| | PNPLA7:DPH7 | √ | | | √ | | √ | | | |
| | SULF2:PRICKLE2 | √ | | | | | | | | |
| | Total | 20 | 1 | 0 | 20 | 15 | 3 | 3 | 17 | 11 |
| MCF-7 Iso-seq | BCAS4:BCAS3 | √ | | | √ | √ | | | | √ |
| | ARFGEF2:SULF2 | | | √ | | | | | | |
| | RPS6KB1:VMP1 | √ | | | √ | √ | | | √ | √ |
| | SMARCA4:CARM1 | | | | | | | | | |
| | SLC25A24:NBPF6 | √ | | | √ | | √ | | √ | |
| | TBL1XR1:RGS17 | √ | | | √ | √ | | | | |

| Dataset | Fusion genes validated | GFHunter | | | LongGF | JAFFAL | | | FusionSeeker | Genion |
|---|---|---|---|---|---|---|---|---|---|---|
| | | RF | SF | PF | | HC | LC | PT | | |
| | RPS6KB1:DIAPH3 | √ | | | √ | | | √ | | |
| | AHCYL1:RAD51C | | | | | | | | | |
| | TXLNG:SYAP1 | | | | | | | | | |
| | MYO6:SENP6 | | | √ | | | | | | |
| | POP1:MATN2 | | | | | | | | | |
| | GATAD2B:NUP210L | | | | | | | | | |
| | ESR1:CCDC170 | √ | | | √ | √ | | | | |
| | DEPDC1B:ELOVL7 | | | | | | | √ | | |
| | ATXN7L3:FAM171A2 | | | | | | | | | |
| | SYTL2:PICALM | √ | | | √ | √ | | | √ | |
| | MYO9B:FCHO1 | | | | | | | | | |
| | PAPOLA:AK7 | | | | √ | | | | | |
| | ATP1A1:ZFP64 | √ | | | | √ | | | | |
| | BCAS3:ATXN7 | | | | | | | | | |
| | BCAS4:ZMYND8 | | | | | | | | | |
| | NAV1:GPR37L1 | | | | | | | | | |
| | NCOA3:SULF2 | | | | | | | | | |
| | PLCG1:TOP1 | | | | | | | | | |
| | PNPLA7:DPH7 | | | | | | | √ | | |
| | SULF2:PRICKLE2 | | | | | | | | | |
| | Total | 8 | 0 | 2 | 8 | 6 | 1 | 3 | 3 | 2 |

RF: “Reliable” fusions; SF: “Suspected” fusions; PF: “Potential” fusions;

HC: “High Confidence”; LC: “Low Confidence”; PT: “Potential Trans-Splicing”.

**Supplementary Table 8. The involved alignments on the uniquely identified two known fusions by GFHunter.**

| Reads id | Alignment type | Flag | Gene | Chrome | Position | Cigar |
|---|---|---|---|---|---|---|
| **HCT-116 dRNA: SPAG9-MBTD1 (HCT-116 cDNA, dRNA)** | | | | | | |
| 39fb3939-7aa6-4588-ad87-e6c19301b7a0 | Genome-based | 2064 | SPAG9 | Chr17 | 51,120,354 | 694H26M2D6M1D6M1I27M2I29M2I7M2D5M1I25M1D20M1I10M3D26M1I76M1D6M1I5M1I17M3D26M1D6M1I7M3I22M1I22M1D11M2I6M1I8M1I10M1D20M1I18M1D37M5H |
| | | 16 | MBTD1 | Chr17 | 51,192,222 | 70S5M1D4M1D8M3I12M5D6M1I6M3D8M501N2M2D5M1D5M2I4M1D26M2D5M1D4M1D12M1I10M1D43M1D32M3I15M1D25M1D4M3D28M411N4M1D7M1I20M1I17M1D13M1D6M4D9M1703N13M2D1M1I41M1D3M1D41M1D4M1I40M6230N1M1I5M1D20M3D27M1I23M1D8M1I10M1D7M3D4M2D4M1D7M512S |
| | Transcriptome-based | 16 | SPAG9 | Chr17 | 13,123,697 | 469S29M4D9M1D43M2D17M1D3M1D11M3D5M1I26M1I2M1I9M2D10M1D5M1D9M2I19M3I31M2D3M1D15M3D6M1I33M1I6M1I20M2I7M2D37M1D6M1D29M1D5M1D8M1D5M1D7M3D9M1I9M6D19M13S |
| | | 2064 | MBTD1 | Chr17 | 13,181,746 | 239H19M3I4M1D12M1D9M2D46M1I4M2I1M1D33M1I6M1D17M6D4M3D9M2I12M1D41M1D11M472H |
| d5a96235-4521-4019-8c60-c67a5e8a4e3d | Genome-based | 16 | SPAG9 | Chr17 | 51,120,356 | 469S29M4D9M1D43M2D17M1D3M1D11M3D5M1I26M1I2M1I9M2D10M1D5M1D9M2I19M3I31M2D3M1D15M3D6M1I33M1I6M1I20M2I7M2D37M1D6M1D29M1D5M1D8M1D5M1D7M3D9M1I9M6D19M13S |
| | | 2064 | MBTD1 | Chr17 | 51,195,008 | 69H12M1D35M3D5M1D15M8D3M1D7M4D10M5D6M2D11M1D1M4D3M2D10M2D1M3D2M2D2M1D15M2D23M3D5M3D4M2D19M3I4M1D12M1D9M2D46M1I4M2I1M1D34M6230N1I5M1D17M6D4M3D9M2I12M1D41M1D7M476H |
| | Transcriptome-based | 16 | SPAG9 | Chr17 | 13,123,697 | 469S29M4D9M1D43M2D17M1D3M1D11M3D5M1I26M1I2M1I9M2D10M1D5M1D9M2I19M3I31M2D3M1D15M3D6M1I33M1I6M1I20M2I7M2D37M1D6M1D29M1D5M1D8M1D5M1D7M3D9M1I9M6D19M13S |
| | | 2064 | MBTD1 | Chr17 | 13,181,746 | 239H19M3I4M1D12M1D9M2D46M1I4M2I1M1D33M1I6M1D17M6D4M3D9M2I12M1D41M1D11M472H |
| **SKBR-3 Iso-seq: CCDC85C-SETD3 (SKBR-3 Iso-seq)** | | | | | | |
| SRR7346977.1263111 | Genome-based | 0 | SETD3 & CCDC85C | Chr14 | 99,397,754 | 53S9M1D403M1D12M1D197M1D94M1D47M1D14M1D119M1D246M1D222M973N161M3965N86M894N167M1144N75M6360N110M1D4M810N59M**122080N**74M36631N113M1645N65M30S |

| Reads id | Alignment type | Flag | Gene | Chrome | Position | Cigar |
|---|---|---|---|---|---|---|
| | Transcriptome-based | 0 | SETD3 | Chr14 | 10,162,583 | 53S9M1D403M1D12M1D197M1D94M1D47M1D14M1D119M1D246M1D821M1D66M279S |
| | | 2048 | CCDC85C | Chr14 | 10,215,243 | 2078H187M95H |
| SRR7346977.1294761 | Genome-based | 16 | SETD3 & CCDC85C | Chr14 | 99,397,748 | 55S274M1D144M1I13M1I325M1D620M973N161M3965N86M894N76M1D51M1D38M1144N75M6360N115M810N59M**122080N**74M43932N146M31S |
| | Transcriptome-based | 16 | SETD3 | Chr14 | 10,162,577 | 55S274M1D144M1I13M1I325M1D943M1D51M1D290M248S |
| | | 2064 | CCDC85C | Chr14 | 10,214,294 | 2094H182M69H |
| SRR7346977.934276 | Genome-based | 0 | SETD3 & CCDC85C | Chr14 | 99,398,416 | 1865S10M1D9M1D5M1I7M1I8M1I8M1I11M1I9M1I4M2I22M1I39M3I4M1I8M1I8M1I14M3I8M5I5M10I8M3I27M1D7M1I16M1I11M1I17M1D6M1I6M1I3M1I4M1D21M3I1M1D7M1I26M4I10M12I6M3D14M1I14M1D12M1D5M1D35M1I21M1I8M3I4M1I5M6I16M1I20M3I3M1I6M2I7M1D6M5I7M6I6M1I3M1I8M1D6M1I8M2I7M320I5219N11M1D5M1I29M2I9M1D9M1I9M3I12M894N33M1D20M5I28M1I58M1I27M1144N36M1D3M2I5M1I5M1I25M6360N18M1I13M1I12M1I13M1I40M1D18M810N3M2I10M1I14M1I11M2D8M1D10M**122080N**5M1D4M2I30M1D11M2D4M1I16M67078N2I2M1I2M1I23M2I4M1I37M59S |
| | Transcriptome-based | 0 | SETD3 | Chr14 | 10,163,221 | 1831S8M3I7M1I2M3I1M1I6M2I10M1D9M1D5M1I7M1I8M1I8M1I11M1I9M1I4M2I22M1I39M1I1M2I3M1I8M1I8M1I14M3I8M5I5M10I8M3I27M1D7M1I16M1I11M1I17M1D6M1I6M1I3M1I4M1D21M3I1M1D7M1I26M1I1M1I2M1I4M1I5M1I1M1I4M5I2M2I14M1I14M1D12M1D5M1D35M1I21M1I8M3I4M1I4M2I2M2I2M1I5M1I8M1I18M1I2M2I3M1I6M2I7M2I7M2I7M6I6M1I3M1I8M1D6M1I8M2I18M1I3M1I3M1I10M1D9M2I5M1I5M1D24M1I5M2I16M3I4M2I8M1I32M1I3M1D10M1I14M3I7M1I13M1I2M1I10M1I11M1D7M2D3M3I8M4I11M2I6M4I11M5I6M1I2M1I12M1I15M1D5M1I29M2I9M1D9M1I9M3I45M1D19M1I4M4I25M1I22M1D9M1I26M1I63M1D3M2I5M1I5M1I43M1I13M1I12M1I13M1I40M1D20M1I2M1I9M1I14M1I11M2D8M1D11M206S |
| | | 2048 | CCDC85C | Chr14 | 10,207,371 | 2379S5M1I8M2I3M1I5M1I1M1I2M1I9M1D2M1D3M1I4M4D3M1I7M1D2M2I19M1I7M1I5M2I3M1I4M1I3M1I3M1D8M5I4M1I7M1I12M1I2M3I6M1I2M1I3M2I9M1I1M6I4M1D3M2I4M1I11M1I5M1D6M1I17M1I2M1D7M6I9M1I3M1I5M2I2 |

| Reads id | Alignment type | Flag | Gene | Chrome | Position | Cigar |
|---|---|---|---|---|---|---|
| | | | | | | M1I6M2I2M3I3M1I5M1D6M1I1M1I4M1I2M2I3M1I2M2I5M1I13M1D2M1I4M3I2M1I3M2I18M2I2M1I1M1I2M1I4M2I15M1I1M1I9M1I6M1I3M1I6M1D10M2I2M2I3M1I3M2D15M1I16M1D6M1I9M2I4M1I2M1D26M1I1M1I7M1I7M1D7M1D6M1I3M1I12M2D3M1D21M1I14M1D16M1D2M2D11M1D11M1D7M2I6M1I3M1I4M1D28M2I8M30I18M3I34M1I9M1I1M1I11M1I1M1I4M1I12M1I6M325S |
| SRR7346977.1135674 | Genome-based | 0 | SETD3 & CCDC85C | Chr14 | 99,405,289 | 19S23M1I10M1D19M1I7M2I10M1I9M1D3M1144N13M1I28M1I10M1D11M2I12M6360N13M3I34M1I8M1I31M2I11M1I2M2I6M1I10M810N22M1I23M2I4M1I4M2I4M3I2M**122080N**43M7I13M1I2M1I3M1I2M1I11M43932N6M2I27M1I15M1I6M2I8M1I11M2I20M1D12M1D12M1D38M1D3M1D60M5I5M2I22M1I16M1I45M34S |
| | Transcriptome-based | 0 | SETD3 | Chr14 | 10,164,269 | 8M1I6M1D2M2I23M1I10M1D19M1I7M1I1M1I9M1I9M1D16M1I28M1I10M1D11M2I13M1I3M1D8M3I34M1I8M1I31M2I11M1I2M2I6M1I32M1I8M1I5M1D10M464S |
| | | 2048 | CCDC85C | Chr14 | 10,214,294 | 377H43M7I13M1I2M1I3M1I2M1I17M2I27M1I15M1I6M2I8M1I11M2I20M1D15M241H |

**Supplementary Table 9. The results of cross validation on five real datasets.**

| | Shared | Doubleton | Singleton | Precision | Recall | F1-score |
|---|---|---|---|---|---|---|
| | | | **HCT116 dRNA** | | | |
| GFHunter[a] | 5 | 0 | 2 | 0.714 | 1.000 | 0.833 |
| LongGF | 5 | 3 | 4 | 0.417 | 1.000 | 0.588 |
| JAFFAL[b] | 5 | 3 | 4 | 0.417 | 1.000 | 0.588 |
| FusionSeeker | 2 | 2 | 42 | 0.043 | 0.400 | 0.078 |
| Genion | 1 | 0 | 38 | 0.026 | 0.200 | 0.045 |
| | | | **SKBR3 Iso-seq** | | | |
| GFHunter[a] | 10 | 1 | 5 | 0.625 | 0.909 | 0.741 |
| LongGF | 10 | 2 | 7 | 0.526 | 0.909 | 0.667 |
| JAFFAL[b] | 11 | 4 | 10 | 0.440 | 1.000 | 0.611 |
| FusionSeeker | 10 | 3 | 35 | 0.208 | 0.909 | 0.339 |
| Genion | 6 | 0 | 43 | 0.122 | 0.545 | 0.200 |
| | | | **MCF7 cDNA** | | | |
| GFHunter[a] | 17 | 3 | 4 | 0.708 | 1.000 | 0.829 |
| LongGF | 16 | 11 | 14 | 0.390 | 0.941 | 0.552 |
| JAFFAL[b] | 15 | 3 | 10 | 0.536 | 0.882 | 0.667 |
| FusionSeeker | 16 | 11 | 31 | 0.276 | 0.941 | 0.427 |
| Genion | 8 | 0 | 6 | 0.571 | 0.471 | 0.516 |
| | | | **MCF7 dRNA** | | | |
| GFHunter[a] | 31 | 2 | 4 | 0.838 | 0.969 | 0.899 |
| LongGF | 32 | 10 | 2 | 0.727 | 1.000 | 0.842 |
| JAFFAL[b] | 26 | 3 | 9 | 0.684 | 0.813 | 0.743 |
| FusionSeeker | 23 | 10 | 29 | 0.371 | 0.719 | 0.489 |
| Genion | 14 | 1 | 17 | 0.438 | 0.438 | 0.438 |
| | | | **MCF7 Iso-seq** | | | |
| GFHunter[a] | 17 | 15 | 42 | 0.230 | 0.810 | 0.358 |
| LongGF | 21 | 64 | 123 | 0.101 | 1.000 | 0.183 |
| JAFFAL[b] | 20 | 48 | 50 | 0.169 | 0.952 | 0.288 |
| Fusionseeker | 9 | 21 | 878 | 0.010 | 0.429 | 0.019 |
| Genion | 2 | 6 | 3 | 0.182 | 0.095 | 0.125 |

a: “Reliable” fusions are used for GFHunter;

b: “High Confidence” and “Low Confidence” fusions are used for JAFFAL.

**Supplementary Table 10. Consistency analysis on three MCF-7 datasets.**

| Harmonic means of consistently reported fusions compared to total reported fusions | | | | | | | |
|---|---|---|---|---|---|---|---|
| | **cDNA+dRNA+Iso-seq** | **cDNA+dRNA** | **cDNA+Iso-seq** | **dRNA+Iso-seq** | **cDNA** | **dRNA** | **Iso-seq** |
| GFHunter RF | 0.133 | 0.492 | 0.163 | 0.144 | - | - | - |
| GFHunter RF+SF | 0.108 | 0.486 | 0.132 | 0.129 | - | - | - |
| LongGF | 0.061 | 0.447 | 0.064 | 0.063 | - | - | - |
| JAFFAL HC+LC | 0.098 | 0.455 | 0.096 | 0.090 | - | - | - |
| FusionSeeker | 0.015 | 0.367 | 0.012 | 0.012 | - | - | - |
| Genion | 0.105 | 0.348 | 0.160 | 0.093 | - | - | - |
| **Count of consistently reported fusions** | | | | | | | |
| | **cDNA+dRNA+Iso-seq** | **cDNA+dRNA** | **cDNA+Iso-seq** | **dRNA+Iso-seq** | **cDNA** | **dRNA** | **Iso-seq** |
| GFHunter RF | 6 | 15 | 8 | 8 | 24 | 37 | 74 |
| GFHunter RF+SF | 6 | 18 | 8 | 9 | 28 | 46 | 93 |
| LongGF | 6 | 19 | 8 | 8 | 41 | 44 | 208 |
| JAFFAL HC+LC | 6 | 15 | 7 | 7 | 28 | 38 | 118 |
| FusionSeeker | 5 | 22 | 6 | 6 | 58 | 62 | 908 |
| Genion | 2 | 8 | 2 | 2 | 14 | 32 | 11 |

RF: “Reliable” fusions; SF: “Suspected” fusions;

HC: “High Confidence”; LC: “Low Confidence”

**Supplementary Table 11. The reported fusions from long-read datasets consistent with short-read datasets.**

| Dataset | Fusion type | Fusion genes reported by short-read | GFHunter | | | LongGF | JAFFAL | | | FusionSeeker | Genion |
|---|---|---|---|---|---|---|---|---|---|---|---|
| | | | RF | SF | PF | | HC | LC | PT | | |
| HCT-116 cDNA | Validated | COMMD10:AP3S1 | √ | | | | | | | √ | √ |
| | | SPAG9:MBTD1 | √ | | | | | | | | |
| | Not yet validated | MAVS:PANK2 | √ | | | | | | | | |
| | | TIPARP:TP53 | | | | | | | | | |
| | Total | 4 | 3 | 0 | 0 | NA | NA | NA | NA | 1 | 1 |
| HCT-116 dRNA | Validated | COMMD10:AP3S1 | √ | | | √ | √ | | | √ | √ |
| | | SPAG9:MBTD1 | √ | | | | | | | | |
| | Not yet validated | MAVS:PANK2 | √ | | | √ | √ | | | | |
| | | TIPARP:TP53 | | | | | | | | | |
| | Total | 4 | 3 | 0 | 0 | 2 | 2 | 0 | 0 | 1 | 1 |
| MCF-7 cDNA | Validated | AHCYL1:RAD51C | √ | | | √ | √ | | | √ | |
| | | AK7:PAPOLA | √ | | | √ | | | √ | | |
| | | ARFGEF2:SULF2 | √ | | | √ | √ | | | √ | |
| | | ATP1A1:ZFP64 | √ | | | √ | √ | | | √ | |
| | | ATXN7L3:FAM171A2 | | | | | | | | | |
| | | BCAS3:BCAS4 | √ | | | √ | √ | | | √ | √ |
| | | BCAS4:ZMYND8 | √ | | | √ | √ | | | √ | √ |
| | | CARM1:SMARCA4 | | | | | | | | | |
| | | CCDC170:ESR1 | | | | | | | | | |
| | | DEPDC1B:ELOVL7 | √ | | | √ | √ | | | √ | |
| | | FCHO1:MYO9B | | | | | | | | | |
| | | GATAD2B:NUP210L | √ | | | √ | √ | | | √ | |
| | | GPR37L1:NAV1 | | | | | | | | | |
| | | MATN2:POP1 | √ | | | | | | √ | | |
| | | MYO6:SENP6 | √ | | | √ | √ | | | √ | |
| | | NBPF6:SLC25A24 | √ | | | √ | | √ | | √ | √ |
| | | PICALM:SYTL2 | | | | | | | | | |
| | | RPS6KB1:VMP1 | √ | | | √ | √ | | | √ | √ |
| | | SYAP1:TXLNG | √ | | | √ | √ | | | √ | √ |
| | Not yet validated | CA4:TANC2 | | | | | | | | | |
| | | EIF4E2:GIGYF2 | √ | | | √ | √ | | | √ | √ |
| | | EMCN:SMARCC1 | √ | | | √ | | | | √ | √ |
| | | GPM6A:SEZ6L2 | √ | | | | √ | | | √ | √ |
| | | TANC2:TLK2 | | | | | | | | | |
| | Total | 24 | 16 | 0 | 0 | 13 | 12 | 1 | 2 | 14 | 8 |
| MCF-7 dRNA | Validated | AHCYL1:RAD51C | √ | | | √ | √ | | | √ | |
| | | AK7:PAPOLA | | √ | | √ | | | √ | √ | |
| | | ARFGEF2:SULF2 | √ | | | √ | √ | | | √ | √ |
| | | ATP1A1:ZFP64 | √ | | | √ | √ | | | √ | √ |
| | | ATXN7L3:FAM171A2 | √ | | | √ | √ | | | √ | √ |
| | | BCAS3:BCAS4 | √ | | | √ | √ | | | √ | √ |
| | | BCAS4:ZMYND8 | √ | | | √ | | √ | | | |
| | | CARM1:SMARCA4 | √ | | | √ | √ | | | √ | |

| Dataset | Fusion type | Fusion genes reported by short-read | GFHunter | | | LongGF | JAFFAL | | | FusionSeeker | Genion |
|---|---|---|---|---|---|---|---|---|---|---|---|
| | | | RF | SF | PF | | HC | LC | PT | | |
| | | CCDC170:ESR1 | | | | | | | | | |
| | | DEPDC1B:ELOVL7 | √ | | | √ | √ | | | √ | √ |
| | | FCHO1:MYO9B | √ | | | √ | √ | | | √ | |
| | | GATAD2B:NUP210L | √ | | | √ | √ | | | √ | √ |
| | | GPR37L1:NAV1 | √ | | | √ | | | √ | | √ |
| | | MATN2:POP1 | | | | | | | | | |
| | | MYO6:SENP6 | √ | | | √ | √ | | | √ | |
| | | NBPF6:SLC25A24 | √ | | | √ | √ | | | √ | √ |
| | | PICALM:SYTL2 | √ | | | √ | √ | | | √ | √ |
| | | RPS6KB1:VMP1 | √ | | | √ | √ | | | √ | √ |
| | | SYAP1:TXLNG | √ | | | √ | √ | | | √ | √ |
| | Not yet validated | CA4:TANC2 | √ | | √ | √ | | | √ | √ | |
| | | EIF4E2:GIGYF2 | | | | | | | √ | | |
| | | EMCN:SMARCC1 | | | | √ | | | | √ | |
| | | GPM6A:SEZ6L2 | | | | | | | | √ | √ |
| | | TANC2:TLK2 | | | | √ | | √ | | √ | |
| | Total | 24 | 17 | 1 | 1 | 20 | 14 | 2 | 4 | 19 | 12 |
| MCF-7 Iso-seq | Validated | AHCYL1:RAD51C | | | | | | | | | |
| | | AK7:PAPOLA | | | | √ | | | | | |
| | | ARFGEF2:SULF2 | | | √ | | | | | | |
| | | ATP1A1:ZFP64 | √ | | | | √ | | | | |
| | | ATXN7L3:FAM171A2 | | | | | | | | | |
| | | BCAS3:BCAS4 | √ | | | √ | √ | | | | √ |
| | | BCAS4:ZMYND8 | | | | | | | | | |
| | | CARM1:SMARCA4 | | | | | | | | | |
| | | CCDC170:ESR1 | √ | | | √ | √ | | | | |
| | | DEPDC1B:ELOVL7 | | | | | | | √ | | |
| | | FCHO1:MYO9B | | | | | | | | | |
| | | GATAD2B:NUP210L | | | | | | | | | |
| | | GPR37L1:NAV1 | | | | | | | | | |
| | | MATN2:POP1 | | | | | | | | | |
| | | MYO6:SENP6 | | | √ | | | | | | |
| | | NBPF6:SLC25A24 | √ | | | √ | | √ | | √ | |
| | | PICALM:SYTL2 | √ | | | √ | √ | | | √ | |
| | | RPS6KB1:VMP1 | √ | | | √ | √ | | | √ | √ |
| | | SYAP1:TXLNG | | | | | | | | | |
| | Not yet validated | CA4:TANC2 | | | | | | | | | |
| | | EIF4E2:GIGYF2 | √ | | | √ | √ | | | | |
| | | EMCN:SMARCC1 | | | | | | | | | |
| | | GPM6A:SEZ6L2 | | | | | | | | | |
| | | TANC2:TLK2 | | | | | | | | | |
| | Total | 24 | 7 | 0 | 2 | 7 | 6 | 1 | 1 | 3 | 2 |

RF: “Reliable” fusions; SF: “Suspected” fusions; PF: “Potential” fusions;

HC: “High Confidence”; LC: “Low Confidence”; PT: “Potential Trans-Splicing”.

**Supplementary Table 12. Time and memory with different total read coverages.**

| | Alignment time/h | Fusion detection time/h | Total time/h | Memory/GB |
|---|---|---|---|---|
| **Total read coverages: 135M** | | | | |
| GFHunter | 3.56 | 0.26 | 3.82 | 10.26 |
| LongGF | 7.54 | 3.63 | 11.17 | 36.28 |
| JAFFAL | 1.90 | 5.61 | 7.51 | 61.04 |
| fusionseeker | 8.09 | 2.71 | 10.80 | 48.49 |
| Genion | 6.80 | 4.06 | 10.86 | 28.83 |
| **Total read coverages: 108M** | | | | |
| GFHunter | 2.86 | 0.17 | 3.03 | 10.97 |
| LongGF | 6.22 | 2.95 | 9.17 | 29.04 |
| JAFFAL | 1.55 | 4.04 | 5.59 | 49.09 |
| fusionseeker | 6.45 | 2.39 | 8.84 | 46.92 |
| Genion | 5.60 | 3.23 | 8.83 | 24.80 |
| **Total read coverages: 81M** | | | | |
| GFHunter | 2.15 | 0.11 | 2.26 | 9.51 |
| LongGF | 4.41 | 2.12 | 6.53 | 21.81 |
| JAFFAL | 1.17 | 2.97 | 4.14 | 37.07 |
| fusionseeker | 4.99 | 1.82 | 6.81 | 46.26 |
| Genion | 4.30 | 2.34 | 6.64 | 22.49 |
| **Total read coverages: 54M** | | | | |
| GFHunter | 1.47 | 0.05 | 1.52 | 9.29 |
| LongGF | 3.21 | 1.43 | 4.64 | 17.87 |
| JAFFAL | 0.83 | 2.00 | 2.83 | 24.69 |
| fusionseeker | 3.36 | 1.23 | 4.59 | 44.70 |
| Genion | 2.77 | 1.53 | 4.30 | 21.61 |
| **Total read coverages: 27M** | | | | |
| GFHunter | 0.76 | 0.03 | 0.79 | 8.77 |
| LongGF | 1.57 | 0.79 | 2.36 | 17.41 |
| JAFFAL | 0.49 | 0.97 | 1.46 | 19.63 |
| fusionseeker | 1.72 | 0.58 | 2.30 | 43.75 |
| Genion | 1.49 | 0.74 | 2.23 | 20.82 |

**Supplementary Table 13. Time and memory with different threads.**

| Detector | Thread | Alignment time (h) | Fusion detection time (h) | Memory (GB) |
|---|---|---|---|---|
| GFHunter | 1 | 3.27 | 0.06 | 11.24 |
| | 2 | 1.66 | 0.06 | 11.70 |
| | 4 | 0.84 | 0.05 | 7.26 |
| | 8 | 0.43 | 0.05 | 6.47 |
| LongGF | 1 | 22.01 | 0.59 | 16.76 |
| | 2 | 11.25 | 0.58 | 17.67 |
| | 4 | 5.63 | 0.58 | 16.63 |
| | 8 | 2.87 | 0.59 | 17.87 |
| JAFFAL | 1 | 2.52 | 0.55 | 18.27 |
| | 2 | 1.29 | 0.41 | 16.69 |
| | 4 | 0.65 | 0.31 | 17.41 |
| | 8 | 0.35 | 0.28 | 20.00 |
| Fusionseeker | 1 | 22.09 | 0.74 | 21.52 |
| | 2 | 11.29 | 0.48 | 22.31 |
| | 4 | 5.65 | 0.37 | 22.47 |
| | 8 | 2.88 | 0.32 | 24.93 |
| Genion | 1 | 21.99 | 0.17 | 16.73 |
| | 2 | 11.25 | 0.17 | 18.27 |
| | 4 | 5.63 | 0.17 | 16.24 |
| | 8 | 2.87 | 0.17 | 18.51 |

**Supplementary Table 14. The availability of the datasets used for benchmark.**

| No. | Dataset | Type | Accession |
|---|---|---|---|
| 1 | gencode.v44.annotation.gtf | Gene annotation | https://ftp.ebi.ac.uk/pub/databases/gencode/Gencode_human/release_44/gencode.v44.annotation.gtf.gz |
| 2 | GCA_000001405.15_GRCh38_no_alt_analysis_set.fna | Reference genome | https://ftp.ncbi.nlm.nih.gov/genomes/all/GCA/000/001/405/GCA_000001405.15_GRCh38/seqs_for_alignment_pipelines.ucsc_ids/GCA_000001405.15_GRCh38_no_alt_analysis_set.fna.gz |
| 3 | HCT-116 short read[1] | Real tumor cell line data | http://sg-nex-data.s3.amazonaws.com/data/sequencing_data_illumina/fastq/SGNex_Hct116_Illumina_replicate3_run1/SGNex_Hct116_Illumina_replicate3_run1_R1.fastq.gz<br>http://sg-nex-data.s3.amazonaws.com/data/sequencing_data_illumina/fastq/SGNex_Hct116_Illumina_replicate3_run1/SGNex_Hct116_Illumina_replicate3_run1_R2.fastq.gz<br>http://sg-nex-data.s3.amazonaws.com/data/sequencing_data_illumina/fastq/SGNex_Hct116_Illumina_replicate4_run1/SGNex_Hct116_Illumina_replicate4_run1_R1.fastq.gz<br>http://sg-nex-data.s3.amazonaws.com/data/sequencing_data_illumina/fastq/SGNex_Hct116_Illumina_replicate4_run1/SGNex_Hct116_Illumina_replicate4_run1_R2.fastq.gz<br>http://sg-nex-data.s3.amazonaws.com/data/sequencing_data_illumina/fastq/SGNex_Hct116_Illumina_replicate5_run1/SGNex_Hct116_Illumina_replicate5_run1_R1.fastq.gz<br>http://sg-nex-data.s3.amazonaws.com/data/sequencing_data_illumina/fastq/SGNex_Hct116_Illumina_replicate5_run1/SGNex_Hct116_Illumina_replicate5_run1_R2.fastq.gz |
| 4 | MCF-7 short read[1] | Real tumor cell line data | http://sg-nex-data.s3.amazonaws.com/data/sequencing_data_illumina/fastq/SGNex_MCF7_Illumina_replicate2_run1/SGNex_MCF7_Illumina_replicate2_run1_R1.fastq.gz<br>http://sg-nex-data.s3.amazonaws.com/data/sequencing_data_illumina/fastq/SGNex_MCF7_Illumina_replicate2_run1/SGNex_MCF7_Illumina_replicate2_run1_R2.fastq.gz<br>http://sg-nex-data.s3.amazonaws.com/data/sequencing_data_illumina/fastq/SGNex_MCF7_Illumina_replicate3_run1/SGNex_MCF7_Illumina_replicate3_run1_R1.fastq.gz<br>http://sg-nex-data.s3.amazonaws.com/data/sequencing_data_illumina/fastq/SGNex_MCF7_Illumina_replicate3_run1/SGNex_MCF7_Illumina_rep |

| No. | Dataset | Type | Accession |
|---|---|---|---|
| | | | licate3_run1_R2.fastq.gz http://sg-nex-data.s3.amazonaws.com/data/sequencing_data_illumina/fastq/SGNex_MCF7_Illumina_replicate4_run1/SGNex_MCF7_Illumina_replicate4_run1_R1.fastq.gz http://sg-nex-data.s3.amazonaws.com/data/sequencing_data_illumina/fastq/SGNex_MCF7_Illumina_replicate4_run1/SGNex_MCF7_Illumina_replicate4_run1_R2.fastq.gz |
| 5 | HG002 Iso-seq | Real non-tumor dataset | https://ftp-trace.ncbi.nlm.nih.gov/ReferenceSamples/giab/data_RNAseq/AshkenazimTrio/HG002_NA24385_son/Baylor_PacBio/reads/m64139_220127_180020.hifi_reads.bam |
| 6 | HG002 cDNA | Real non-tumor dataset | https://s3.amazonaws.com/gtl-public-data/giab/bams/cDNA/05_09_23_R941_GIAB_cDNA_PCS111_NA24385_Guppy_6.4.6_sup.pass.fastq.gz.hg38.bam https://s3.amazonaws.com/gtl-public-data/giab/bams/cDNA/05_09_23_R941_GIAB_cDNA_PCS111_NA26105_Guppy_6.4.6_sup.pass.fastq.gz.hg38.bam https://s3.amazonaws.com/gtl-public-data/giab/bams/cDNA/05_09_23_R941_GIAB_cDNA_PCS111_NA27730_Guppy_6.4.6_sup.pass.fastq.gz.hg38.bam |
| 7 | HG002 dRNA | Real non-tumor dataset | https://s3.amazonaws.com/gtl-public-data/giab/bams/dRNA/03_30_23_R941_DRS_NA24385_dRNA_Guppy_6.4.6_rna_hac_prom.pass.NoU.fastq.gz.hg38.bam |
| 8 | HCT-116 cDNA[1] | Real tumor cell line data | http://sg-nex-data.s3.amazonaws.com/data/sequencing_data_ont/fastq/SGNex_Hct116_cDNA_replicate1_run6/SGNex_Hct116_cDNA_replicate1_run6.fastq.gz http://sg-nex-data.s3.amazonaws.com/data/sequencing_data_ont/fastq/SGNex_Hct116_cDNA_replicate3_run3/SGNex_Hct116_cDNA_replicate3_run3.fastq.gz http://sg-nex-data.s3.amazonaws.com/data/sequencing_data_ont/fastq/SGNex_Hct116_cDNA_replicate3_run5/SGNex_Hct116_cDNA_replicate3_run5.fastq.gz http://sg-nex-data.s3.amazonaws.com/data/sequencing_data_ont/fastq/SGNex_Hct116_cDNA_replicate4_run2/SGNex_Hct116_cDNA_replicate4_run2.fastq.gz http://sg-nex-data.s3.amazonaws.com/data/sequencing_data_ont/fastq/SGNex_Hct116_cDNA_replicate4_run4/SGNex_Hct116_cDNA_replicate4_run4.fastq.gz |

| No. | Dataset | Type | Accession |
|---|---|---|---|
| | | | http://sg-nex-data.s3.amazonaws.com/data/sequencing_data_ont/fastq/SGNex_Hct116_cDNA_replicate4_run5/SGNex_Hct116_cDNA_replicate4_run5.fastq.gz |
| 9 | HCT-116 dRNA[1] | Real tumor cell line data | http://sg-nex-data.s3.amazonaws.com/data/sequencing_data_ont/fastq/SGNex_Hct116_directRNA_replicate1_run1/SGNex_Hct116_directRNA_replicate1_run1.fastq.gz<br>http://sg-nex-data.s3.amazonaws.com/data/sequencing_data_ont/fastq/SGNex_Hct116_directRNA_replicate1_run2/SGNex_Hct116_directRNA_replicate1_run2.fastq.gz<br>http://sg-nex-data.s3.amazonaws.com/data/sequencing_data_ont/fastq/SGNex_Hct116_directRNA_replicate1_run3/SGNex_Hct116_directRNA_replicate1_run3.fastq.gz<br>http://sg-nex-data.s3.amazonaws.com/data/sequencing_data_ont/fastq/SGNex_Hct116_directRNA_replicate2_run1/SGNex_Hct116_directRNA_replicate2_run1.fastq.gz<br>http://sg-nex-data.s3.amazonaws.com/data/sequencing_data_ont/fastq/SGNex_Hct116_directRNA_replicate2_run2/SGNex_Hct116_directRNA_replicate2_run2.fastq.gz<br>http://sg-nex-data.s3.amazonaws.com/data/sequencing_data_ont/fastq/SGNex_Hct116_directRNA_replicate2_run3/SGNex_Hct116_directRNA_replicate2_run3.fastq.gz<br>http://sg-nex-data.s3.amazonaws.com/data/sequencing_data_ont/fastq/SGNex_Hct116_directRNA_replicate2_run4/SGNex_Hct116_directRNA_replicate2_run4.fastq.gz<br>http://sg-nex-data.s3.amazonaws.com/data/sequencing_data_ont/fastq/SGNex_Hct116_directRNA_replicate2_run5/SGNex_Hct116_directRNA_replicate2_run5.fastq.gz<br>http://sg-nex-data.s3.amazonaws.com/data/sequencing_data_ont/fastq/SGNex_Hct116_directRNA_replicate2_run6/SGNex_Hct116_directRNA_replicate2_run6.fastq.gz<br>http://sg-nex-data.s3.amazonaws.com/data/sequencing_data_ont/fastq/SGNex_Hct116_directRNA_replicate3_run1/SGNex_Hct116_directRNA_replicate3_run1.fastq.gz<br>http://sg-nex- |

| No. | Dataset | Type | Accession |
|---|---|---|---|
| | | | data.s3.amazonaws.com/data/sequencing_data_ont/fastq/SGNex_Hct116_directRNA_replicate3_run4/SGNex_Hct116_directRNA_replicate3_run4.fastq.gz<br>http://sg-nex-data.s3.amazonaws.com/data/sequencing_data_ont/fastq/SGNex_Hct116_directRNA_replicate4_run3/SGNex_Hct116_directRNA_replicate4_run3.fastq.gz<br>http://sg-nex-data.s3.amazonaws.com/data/sequencing_data_ont/fastq/SGNex_Hct116_directRNA_replicate6_run1/SGNex_Hct116_directRNA_replicate6_run1.fastq.gz |
| 10 | SKBR-3 Iso-seq[2] | Real tumor cell line data | SRP150606 |
| 11 | MCF-7 cDNA[1] | Real tumor cell line data | http://sg-nex-data.s3.amazonaws.com/data/sequencing_data_ont/fastq/SGNex_MCF7_cDNA_replicate1_run3/SGNex_MCF7_cDNA_replicate1_run3.fastq.gz<br>http://sg-nex-data.s3.amazonaws.com/data/sequencing_data_ont/fastq/SGNex_MCF7_cDNAStranded_replicate2_run1/SGNex_MCF7_cDNAStranded_replicate2_run1.fastq.gz<br>http://sg-nex-data.s3.amazonaws.com/data/sequencing_data_ont/fastq/SGNex_MCF7_cDNAStranded_replicate3_run2/SGNex_MCF7_cDNAStranded_replicate3_run2.fastq.gz |
| 12 | MCF-7 dRNA[1] | Real tumor cell line data | http://sg-nex-data.s3.amazonaws.com/data/sequencing_data_ont/fastq/SGNex_MCF7_directRNA_replicate1_run1/SGNex_MCF7_directRNA_replicate1_run1.fastq.gz<br>http://sg-nex-data.s3.amazonaws.com/data/sequencing_data_ont/fastq/SGNex_MCF7_directRNA_replicate2_run2/SGNex_MCF7_directRNA_replicate2_run2.fastq.gz<br>http://sg-nex-data.s3.amazonaws.com/data/sequencing_data_ont/fastq/SGNex_MCF7_directRNA_replicate2_run3/SGNex_MCF7_directRNA_replicate2_run3.fastq.gz<br>http://sg-nex-data.s3.amazonaws.com/data/sequencing_data_ont/fastq/SGNex_MCF7_directRNA_replicate3_run1/SGNex_MCF7_directRNA_replicate3_run1.fastq.gz<br>http://sg-nex-data.s3.amazonaws.com/data/sequencing_data_ont/fastq/SGNex_MCF7_directRNA_replicate4_run1/SGNex_MCF7_directRNA_replicate4_run1.fastq.gz |
| 13 | MCF-7 Iso-seq[3] | Real tumor cell line data | SRP055913 |

**Supplementary Table 15. All the tools used in this study**

| Tools | Version | URL | Category |
|---|---|---|---|
| Minimap2 | 2.22(r1101) | https://github.com/lh3/minimap2 | Long read sequence aligner |
| abPOA | v1.5.2 | https://github.com/yangao07/abPOA | Multiple sequence aligner |
| LongGF | v0.1.2 | https://github.com/WGLab/LongGF | Long read fusion detection tools |
| JAFFAL | Version 2.3 | https://github.com/Oshlack/JAFFA | Long read fusion detection tools |
| FusionSeeker | v1.0.1 | https://github.com/Maggi-Chen/FusionSeeker | Long read fusion detection tools |
| Genion | 1.1.1 | https://github.com/vpc-ccg/genion | Long read fusion detection tools |
| JAFFA | Version 2.3 | https://github.com/Oshlack/JAFFA | Short read fusion detection tools |
| STAR-Fusion | v1.13.0 | https://github.com/STAR-Fusion/STAR-Fusion | Short read fusion detection tools |
| PBSim3 | v3.0.4 | https://github.com/yukiteruono/pbsim3 | Long read sequencing simulator |
| Samtools[4] | 1.13 | https://github.com/samtools/samtools | SAM processing tool |

### Supplementary Notes

#### Implementation of simulation

Here's a streamlined and technically precise description of the simulated data construction process for gene fusion detection:

Step 1: Access and read the gene annotation file to document the start and end positions of the exons for each transcript. Utilize these positional details to locate and extract the corresponding exon sequences from the reference genome. Then, refine this dataset by filtering out transcripts, retaining only those that are classified as protein-coding and comprise three or more exons.

Step 2: Randomly select one transcript from each of 1,000 distinct genes. Divide these selected transcripts into two groups. Given that these transcripts originate from different genes, they can be freely combined to facilitate the generation of fusion transcripts.

Step 3: Select transcript pairs from the two groups and generate 500 fusion transcripts for each pair. The following steps outline the processing for each pair of transcripts: 1) randomly choose non-boundary exons from each transcript to serve as breakpoints; 2) establish the fusion pattern based on chromosome strand information (refer to Fig. 2a); 3) keep the exons consistent with the determined fusion pattern; 4) connect the retained exons and randomly trim between 0 to 20 base pairs at the breakpoints to enhance data complexity; 5) merge the retained sequences from the two transcripts to form the complete fusion transcript. Note that the number of fusion transcripts produced for each transcript pair is contingent upon the simulated depth.

Step 4: Trim both ends of each fusion transcript sequence generated in the previous step, removing anywhere from 0 up to half the length of the remaining sequence. After trimming, retain only those fusion transcripts that are 100 base pairs or longer in length.

Step 5: Utilize the processed fusion transcripts as input and employ pbsim3's template-based read generation function to introduce stochastic variability into the transcripts, and then generate simulated reads.

**Positive simulations on ONT with pbsim3 and samtools**

```
pbsim --strategy templ \
      --method errhmm \
      --errhmm ERRHMM-ONT-HQ.model \
      --accuracy-mean 0.85 \
      --prefix GF \
      --id-prefix GF \
      --template simulation.fasta

samtools fasta "GF.fastq" > "ont_simulation_depth.fasta"

rm -rf *.bam* *.sam *.txt *.gz *.maf *.fastq
```

**Positive simulations on PacBio with pbsim3 and samtools**

```
pbsim --strategy templ \
```

```
		--method errhmm \
		--errhmm ERRHMM-SEQUEL.model \
		--accuracy-mean 0.85 \
		--prefix GF \
		--id-prefix GF \
		--template simulation.fasta

	samtools fasta "GF.fastq" > "ont_simulation_depth.fasta"

	rm -rf *.bam* *.sam *.txt *.gz *.maf *.fastq
```

**Negative simulations on ONT with pbsim3 and samtools**

```
	for ((i = 1; i <= 25; i++ ))
	do
	num="_$i"
	name=$type$num

	pbsim --strategy wgs \
		--method qshmm \
		--qshmm QSHMM-ONT.model \
		--depth 10 \
		--prefix $name \
		--id-prefix "Normal""$i"_ \
		--genome "gencode.v44.transcripts.length_over_100.$i.fa"

	rm -rf *.maf *.ref
	cat Normal*.fastq &gt; "all_$i.fastq"
	rm Normal*.fastq
	samtools fasta "all_$i.fastq" &gt; "all_$i.fasta"
	rm "all_$i.fastq"
	done

	cat all_*.fasta &gt; ../Normal_ONT_10X.fasta
	rm all_*.fasta
```

**Negative simulations on PacBio with pbsim3 and samtools**

```
	for ((i = 1; i <= 25; i++ ))
	do
	num="_$i"
	name=$type$num

	pbsim --strategy wgs \
		--method qshmm \
		--qshmm QSHMM-RSII.model \
```

```
        --depth 10 \
        --prefix $name \
        --id-prefix "Normal""$i"_ \
        --genome "gencode.v44.transcripts.length_over_100.$i.fa"

rm -rf *.maf *.ref
cat Normal*.fastq &gt; "all_$i.fastq"
rm Normal*.fastq
samtools fasta "all_$i.fastq" &gt; "all_$i.fasta"
rm "all_$i.fastq"
done

cat all_*.fasta &gt; ../Normal_PacBio_10X.fasta
rm all_*.fasta
```

#### The command lines for fusion identification

**GFHunter**

```
GFHunter detect \
        -o name \
        -t 32 \
        -M \
        XXX.fasta \
        ./index/
```

**LongGF**

```
minimap2 -ax splice \
        -t 32 \
        ref_index.mmi \
        XXX.fasta \
        -o .sam
samtools view -b .sam -o .bam
LongGF .bam \
      gencode.v44.annotation.gtf \
      100 30 100 \
      > name.log
```

**JAFFAL**

```
JAFFA-master/tools/bin/bpipe run -n 32\
        JAFFA-master/JAFFAL.groovy \
        XXX.fasta
```

**FusionSeeker**

```
minimap2 -ax splice \
        -t 32 \
```

```
            ref_index.mmi \
            XXX.fasta \
            -o .sam
    samtools sort .sam -o .bam
    samtools index .bam
    FusionSeeker --bam .bam \
                -o FusionSeeker_out/ \
                --gtf gencode.v44.annotation.gtf \
                --ref GCA_000001405.15_GRCh38_no_alt_analysis_set.fa \
                --thread 32
```

**Genion**

```
    minimap2 -x splice \
            -c \
            -t 32 \
            ref_index.mmi \
            XXX.fasta \
            -o.paf
    genion -i XXX.fasta \
            --gtf small_example/Homo_sapiens.GRCh38.97.gtf \
            --gpaf .paf \
            -s /small_example/cdna.self.tsv \
            -d /small_example/genomicSuperDups.txt \
            -o name.tsv
```

**JAFFA**

```
    JAFFA-master/tools/bin/bpipe run -n 30\
            JAFFA-master/JAFFA_direct.groovy \
            R1.fasta \
            R2.fasta
```

**STAR-Fusion**

```
    singularity exec \
        -e \
        -B `pwd` \
        -B                    starfusion/GRCh38_gencode_v44_CTAT_lib_Oct292023.plug-n-
    play/ctat_genome_lib_build_dir \
        /home/downloads/images/sif/star-fusion.v1.13.0.simg \
        STAR-Fusion \
        --left_fq R1.fastq.gz \
        --right_fq R2.fastq.gz \
        --genome_lib_dir    ngsgf/starfusion/GRCh38_gencode_v44_CTAT_lib_Oct292023.plug-n-
    play/ctat_genome_lib_build_dir \
        -O StarFusionOut \
```

```
--FusionInspector validate \
--examine_coding_effect \
--denovo_reconstruct
```

##### The command lines for evaluation

All the detailed command lines are in-house scripts that can be assessed at GitHub: https://github.com/luzhenhao-HIT/GFHunter.
